# Higher-order Residue Interactions Encode Diversified Physical Properties of Biomolecular Condensates

**DOI:** 10.1101/2025.09.13.675986

**Authors:** Le Chen, Xiangyu Wu, Guorong Hu, Xiangze Zeng, Jingyuan Li

## Abstract

Biomolecular condensates often formed by proteins containing extended intrinsic disordered regions (IDPs) can coordinate versatile cellular processes. The biological functions of condensates are attributed to a variety of physical properties, including viscoelasticity, interfacial tension and saturation concentration. Comprehensive understanding of biological functions of protein condensates and the corresponding sequence encoding requires unified interpretation of the diverse physical properties by a single model. Here we report the diversified physical properties of condensate and the interactions between constituent IDPs can be interpreted on the basis of higher-order residue interactions. These findings are based on μs-scale atomistic molecular dynamics simulations of three homogeneous IDP condensates (FUS PLD, ARF6 PLD, LAF-1 RGG) combined with the reported simulation trajectories of two heterogeneous condensates. In all interacting IDPs, there are residue interactions between finite motifs (3∼5 residues) that organize in a highly cooperative way: residues form multivalent interactions and their remaking is localized. The sequence features of higher-order interactions (HOIs) are identified: tyrosine, arginine, glutamine and aspartic acid are essential, and their consecutive sequence position is also important. The interaction between corresponding motifs is highly sustained and dominates the cohesive energy of IDP. IDP condensate can be modeled as a HOI-crosslinked network. This model can successfully reproduce all reported experimental results of condensate viscoelasticity, interfacial tension and saturation concentration, and expand our insights into the molecular grammar of IDP interactions.

**Significance Statement:** The biological functions of biomolecular condensates involve various physical properties of condensate. A unified model that describes all these physical properties is essential for the understanding of condensate functions. Our research reveals the prevalence of higher-order residue interactions in all IDP condensates we considered, which leads to extremely sustained interaction between corresponding residue motifs. Accordingly, we propose a physically crosslinked network based on higher-order residue interactions as a simplified model for condensate. This model can quantitively reproduce all reported experimental results of condensate physical properties, including viscoelasticity, interfacial tension and saturation concentration.

## Introduction

Biomolecular condensates are membraneless bodies that usually form through liquid-liquid phase separation (LLPS) of proteins containing extended intrinsic disordered regions (IDPs) and other biomolecules (1-8). Condensates can coordinate versatile cellular processes, e.g., gene regulation, signaling and stress response (2, 5, 9, 10), and each process involves various activities of condensate like storage of biomolecules and wetting with membrane structures (9, 11-16). These activities are related to a spectrum of condensate physical properties including viscoelasticity, interfacial tension and saturation concentration (11, 14, 17-22). Integrative comprehension of the mechanism underlying these physical properties, i.e., the unified interpretation, is essential for the better understanding of the biological functions of condensate (23-27).

There have been extensive studies about the diverse physical properties of condensate. Condensates usually exhibit high viscoelasticity and low interfacial tension (10, 28, 29). Both exhibit significant deviation from standard oil droplets like long-chain alkanes (29-31): the viscosities of condensate are at least 100 times higher, whereas the interfacial tension coefficients are at least 100 times lower (Fig. S1). Thus, there is remarkable decoupling between these two physical properties. Experimental results of condensate viscosity are usually reproduced based on Rouse model of polymer solutions (23, 32, 33). While the viscoelasticity of condensate can be characterized by Maxwell model (30, 34, 35), its calculation requires the parameter from experimental results (32, 33). The saturation concentration of the formation of some condensates can be calculated based on stickers-and-spacers model (36-38). The study on condensate interfacial tension is still limited. To date, only a fraction of physical properties can be directly reproduced by these models and these models interpret the molecular IDP interactions (their stability and dynamics) in diversified ways (17, 23, 33, 38). Integrative comprehension of these physical properties by a single model is essential for the better understanding of the biological functions of condensate, and consensual interpretation of IDP interactions is essential for this unified model of condensate (17).

The protein interactions in condensates have been extensively studied in the experiments. IDP interactions are conceived as weak, multivalent interactions (4, 39, 40). Specific residues, such as tyrosine and arginine, are recognized to make important contributions to IDP interactions and intensively discussed in stickers-and-spacers model (33, 35-38, 41, 42). On the other hand, there is growing evidence about the contributions of other residues like glutamine and aspartic acid, as well as the importance of residue patterning (37, 43-50). Comprehensive investigation of IDP interactions at the residue level, especially the description of the involvement of various residues as well as their cooperativity, can advance our understanding about sequence feature of IDP interaction (i.e., molecular grammar) as well as the physical properties of condensate (44, 51-54).

Atomistic molecular dynamics (MD) simulations of IDP condensate can provide detailed information about IDP interactions (23, 43, 44, 55-58). For example, dynamic IDP interactions are depicted as the making and breaking of residue contacts (23, 55). And the probability of the involvement of residues to IDP interactions as well as all pair-wise residue contacts are scrutinized (43, 44, 57). In this study, we conducted μs-scale atomistic molecular dynamics simulations of three homogeneous IDP condensates (FUS PLD, ARF6 PLD, LAF-1 RGG) combined with the reported simulation trajectories of two heterogeneous condensates (H1-ProTα, protamine-ProTα) (23, 55) to systematically study the IDP interactions. In all these condensates, there is a fraction of (∼30%) residue contacts with the lifetime about ten times longer than other residue contacts. These long-lived contacts are concentrated on the finite residue motifs (3∼5 residues). Notably, the lifetimes of interaction between these residue motifs reach thousands of nanoseconds: three orders of magnitude longer than most residue contacts. Such sustained interaction between residue motifs is denoted as *motif interaction*. Within these motifs, residue interactions organize in a highly cooperative way: most residues have multivalent interactions and their reorganizations are localized. The sequence features of such higher-order interactions (HOIs) are identified: tyrosine, arginine, glutamine and aspartic acid are important. Besides, the consecutive sequence position of these important residues is also essential. HOIs between finite motifs essentially dominate IDP interactions and the corresponding cohesive energy of IDPs. Accordingly, IDP condensate can be conceived as HOI-crosslinked network. Our proposed model can successfully reproduce the experimental results of condensate viscoelasticity, interfacial tension and saturation concentration, providing a unified and quantitative interpretation of seemingly diversified physical properties of condensate.

## Results

### *Motif interaction* identified in all condensates we considered

Three IDP condensates were constructed in this work, including FUS PLD enriched in aromatic residues, ARF6 PLD enriched in polar residues and LAF-1 RGG enriched in charged residues (*SI Appendix*, Fig. S2). The initial configurations for atomistic simulations of these systems were obtained from a coarse-grained (CG) simulation based on the procedure described before (57). Three 3-μs atomistic MD simulations were performed for FUS PLD condensate (Fig. 1A) and one 3-μs simulation for both ARF6 PLD and LAF-1 RGG condensate (Fig. S3 and S4). Further simulation details are provided in the *SI Appendix*.

**Fig. 1.**
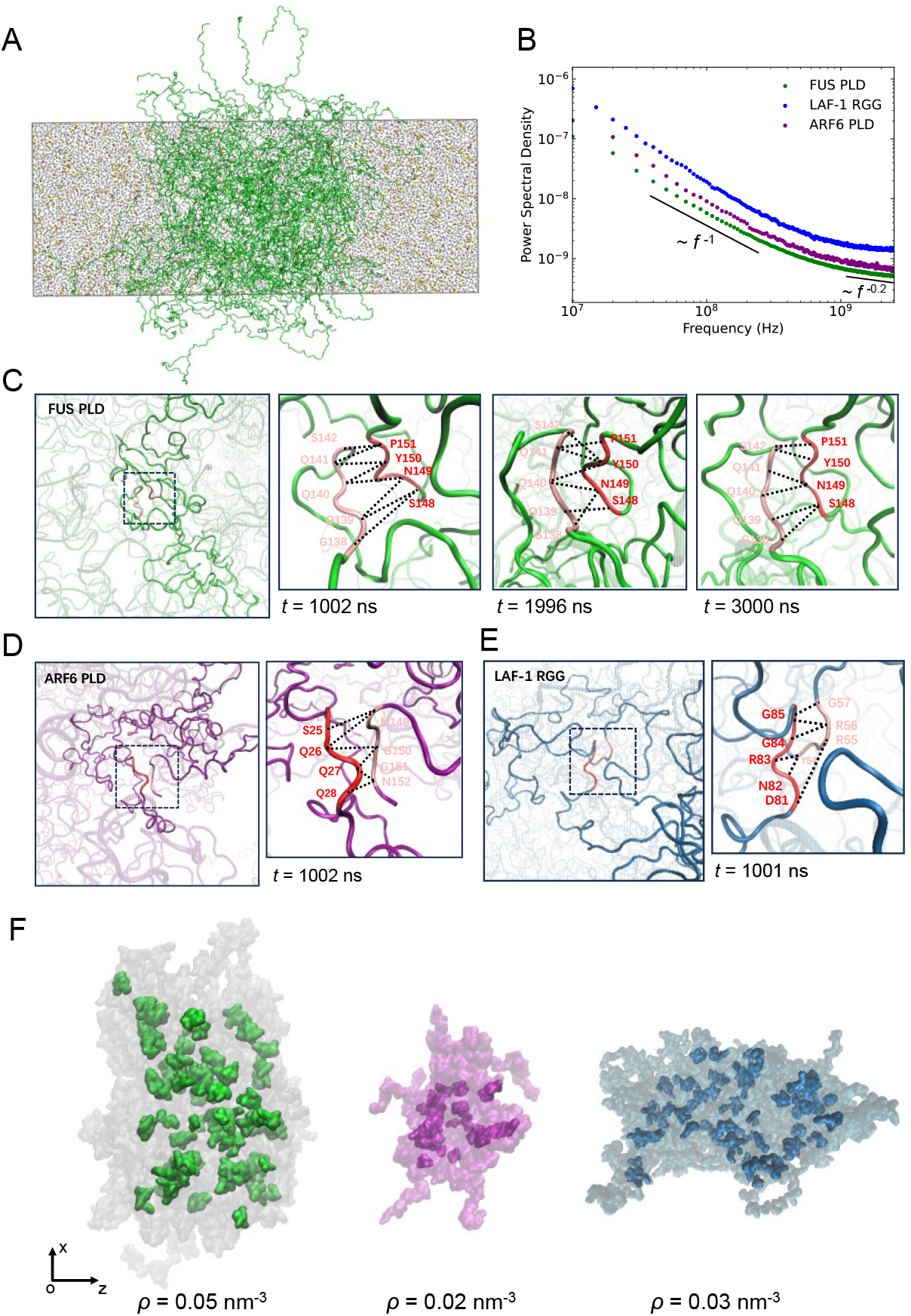
*Motif interaction* identified in three condensates. (A) Representative snapshot of atomistic explicit-solvent simulation of FUS PLD condensate. Proteins are shown explicitly in green strips, with water colored in gray, and Na+ and Cl-ions in red and yellow, respectively. (B) Power spectral density (PSD) of the evolution of the number of residue contacts between IDPs (*N*_*C*_ (*t*)) in FUS PLD (green), ARF6 PLD (purple) and LAF-1 RGG (blue) condensate. (C, D and E) Representative configurations of *motif interaction*s within FUS PLD (C), ARF6 PLD (D) and LAF-1 RGG (E) condensates and localized reorganization of residue interaction in *motif interaction*s. Two residue motifs are colored in red and pink, and the residue interactions are highlighted in black dashed lines. (F) Spatial distributions of *motif interaction*s within FUS PLD (left panel), ARF6 PLD (middle panel) and LAF-1 RGG (right panel) condensates at *t* ≈ 2000 ns, along with the corresponding density (*ρ*) of *motif interaction*s.

The intermolecular interactions between IDPs in three condensates were analyzed. The evolution of the number of residue contacts (under the contact definition of a 0.4 nm distance between heavy atoms) between IDPs, *N*_*C*_(*t*), is presented in Fig. S5. *N*_*C*_(*t*) continuously undergoes fluctuation, indicating dynamic making and breaking of residue contacts. Temporal correlation (56) can be depicted by the power spectral density (PSD) (59) of *N*_*C*_(*t*) (Fig. 1B). For all three systems, the PSD exhibits a plateau above a frequency of ∼10^9^ Hz. Notably, at frequencies ∼10^8^ Hz, PSD exhibits 1/*f* noise with a power-law exponent of ∼-1.2. Hence, there are residue contacts with relatively long lifetimes.

The survival probability (60) of residue contacts between IDPs *S*(*t*) (the ratio of contacts that survive until time *t*) is shown in Fig. S6. We notice that the survival probability can be described as a double-exponential function (colored in red):

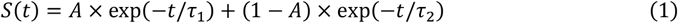

where τ_1_ and τ_2_ are the time constants for the slow and fast decay modes and *A* and (1 − A) are the corresponding weighting factors. Three IDP condensate share similar decay modes with τ_1_ = 24.25 ± 0.52 ns, τ_2_ = 1.07 ± 0.02 ns (FUS PLD); τ_1_ = 18.79 ± 0.37 ns, τ_2_ = 0.79 ± 0.02 ns (ARF6 PLD); τ_1_ = 16.80 ± 0.45 ns, τ_2_ = 0.99 ± 0.03 ns (LAF-1 RGG). And the partition coefficient *A* = 0.346 ± 0.004 (FUS PLD), *A* = 0.322 ± 0.004 (ARF6 PLD), *A* = 0.313 ± 0.003 (LAF-1 RGG). In short, there is a fraction of long-lived residue contacts in all three IDP condensates.

The prolonged residue contacts were then identified by means of the threshold lifetime (60): the weighted mean of time constants 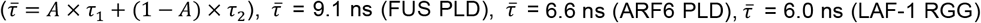 Fig. 1C shows a representative configuration of prolonged residue contacts between an interacting FUS PLD pair. At *t* = 1002 ns, there are eight prolonged residue contacts (highlighted in black dashed lines). Interestingly, they involve residue motifs from two IDPs: residue motif 138-142 of IDP-1 and residue motif 148-151 of IDP-2. Meanwhile, there are normal residue contacts (∼1 ns) in the vicinity of these localized prolong contacts (Fig. S8). For example, Q145 of IDP-1 can contact with Q160 of IDP-2. And this contact quickly breaks at *t* = 1007.4 ns. Subsequently, Q145 can form transient contact with S163 of IDP-2.

At *t* ≈ 2000 ns, there are seven prolonged residue contacts (Fig.1 C). Interestingly, they still locate between same residue motifs (i.e. 138-142 and 148-151). And some contacts have underwent breaking and remaking. For example, S142 no longer contacts with P151 and instead forms contact with Y150. At the end of the simulation (i.e., *t* ≈ 3000 ns), these two residue motifs still keep contact and there are six prolonged residue contacts (Fig. 1C). Hence, residue motifs 138-142 and 148-151 keep contact for over 2000 ns (Fig. S7) and the lifetime is three orders of magnitude longer than most residue contacts (∼ns)! Taken together, the residue interactions across these two residue motifs are highly coupled: the residues can form multivalent interaction with the other residue motif and the remaking of residue interactions is very localized. Highly cooperative residue interactions should contribute to the extremely sustained interaction between the corresponding motifs (with the lifetime more than 1000 ns, denoted as *motif interaction*).

For FUS PLD condensate, there are 626 interacting IDP pairs (with the threshold value of residue contacts of 30) observed in the combined 9-μs trajectories (Table S3). Notably, *motif interaction* occurs between all these interacting IDP pairs (Fig. 1F, left panel; Fig. S12). And the number of *motif interaction* between each interacting IDP pair is 1.68. There are 7.69 residue contacts between two motifs with the length of 3.79. It is much more than the number of residue contacts between two random residue segments with the same length (one interacting FUS PLD pair forms 36.38 residue contacts, corresponding to <1 contact between two 4-residue segment). And there are 2.54 hydrogen bonds within *motif interaction*. We further discussed the lifetime of such cooperative *motif interaction*. According to Arrhenius’ law (61), the lifetime of an isolated residue interaction is given by τ_1_ = *a* exp(Δ*G*_1_/*k*_*B*_*T*), where Δ*G*_1_ is the free energy of the isolate residue interaction. And the lifetime of *motif interaction* involving cooperative residue contacts can be expressed as τ_m_ = *a* exp(Δ*G*_m_/*k*_*B*_*T*), where Δ*G*_m_ is the free energy of *motif interaction*. As mentioned above, the residues can form sufficient interactions and their reorganizations are localized within *motif interaction*. The difference between Δ*G*_m_ and Δ*G*_1_ can be manifested as the enthalpic gains solely from the additional interactions (35, 62, 63), i.e., Δ*G*_m_ = Δ*G*_1_ + ΔΔ*H*_m_. And the enhancement in lifetime of *motif interaction* with respect to the isolated residue interaction θ = τ_m_/τ_1_ = exp (ΔΔ*H*_m_/*k*_*B*_*T*). On average, *motif interaction* forms 1.54 additional hydrogen bonds. The average strength of a hydrogen bond in protein is ∼4*k*_B_T (64-66) and the contribution of additional hydrogen bond is ∼6.16*k*_B_T. Combined with the other contribution, such as π-interaction, ΔΔ*H*_m_ should be beyond 7*k*_B_T and θ is >1000. And it is consistent with our simulation results.

Taken together, the residues between these finite motifs form sufficient interactions and their reorganizations are localized. Moreover, the contribution of such cooperative interactions to their lifetime can be attributed to non-additive effect of residue interaction and be interpreted as enthalpic gains. In short, these highly cooperative residue interactions can be conceived as higher-order interactions (HOIs) between corresponding residue motifs.

We also studied the location of prolonged residue contacts in ARF6 PLD (Fig. 1D) and LAF-1 RGG (Fig. 1E) condensates. Interestingly, prolonged residue contacts can also be attributed to the interaction between the residue motifs from different IDPs (i.e., 25-28 and 149-152 in ARF6 PLD, 55-57 and 81-85 in LAF-1 RGG). The residue interactions are also organized in a highly cooperative way, and the lifetime of *motif interaction* in these two condensates also exceeds 1000 ns (Fig. S7). Moreover, *motif interaction* also exists between all interacting IDP pairs of both condensates (Fig. 1F, middle and right panel; Fig. S12), with average number of 1.19 (ARF6 PLD) and 1.73 (LAF-1 RGG). It is important to note that, sustained *motif interaction* can be also identified in the reported simulation of two hybrid condensates, i.e., H1-ProTα condensate and protamine-ProTα condensate (Fig. S9). And the average number of *motif interaction* between interacting IDP pairs is 1.36 (H1-ProTα) and 1.40 (protamine-ProTα). Taken together, such sustained interaction mode, i.e., *motif interaction*, is commonly present in a variety of condensates we studied. And these *motif interaction*s are organized in a highly cooperative way and can be conceived as the higher-order interaction between finite motifs.

### Distribution of *motif interaction*s in the condensate

The spatial distributions of *motif interaction*s within three homogenous condensates were studied in a systematic way. Representative configurations of *motif interaction*s are shown in Fig. 1F. *Motif interaction*s scatter throughout the condensate. Yet their densities are quite low: 0.05 nm^-3^ (FUS PLD), 0.02 nm^-3^ (ARF6 PLD) and 0.03 nm^-3^ (LAF-1 RGG), at least two orders of magnitude lower than residues (3∼7 nm^-3^).

It should be noted that spatial arrangement of *motif interaction* is quite stable (Fig. 2A and Fig. S10). Only a small fraction of *motif interaction*s undergoes slight slippage (black arrows). Therefore, steady arrangement of sustained *motif interaction* results in the resilient configuration of condensate even though most residue contacts are highly dynamic. Besides, the interaction energy associated with *motif interaction* accounts for >50% of the total interaction energy between IDPs (*SI Appendix*, Table S4), which is largely attributed to the higher-order residue interaction between corresponding motifs. Thus, HOI-mediated *motif interaction* dominates the protein interaction in condensates. Taken together, the condensate can be conceived as physically crosslinked network (4, 67-69) mediated by HOI-mediated *motif interaction*, i.e., HOI-crosslinked network.

**Fig. 2.**
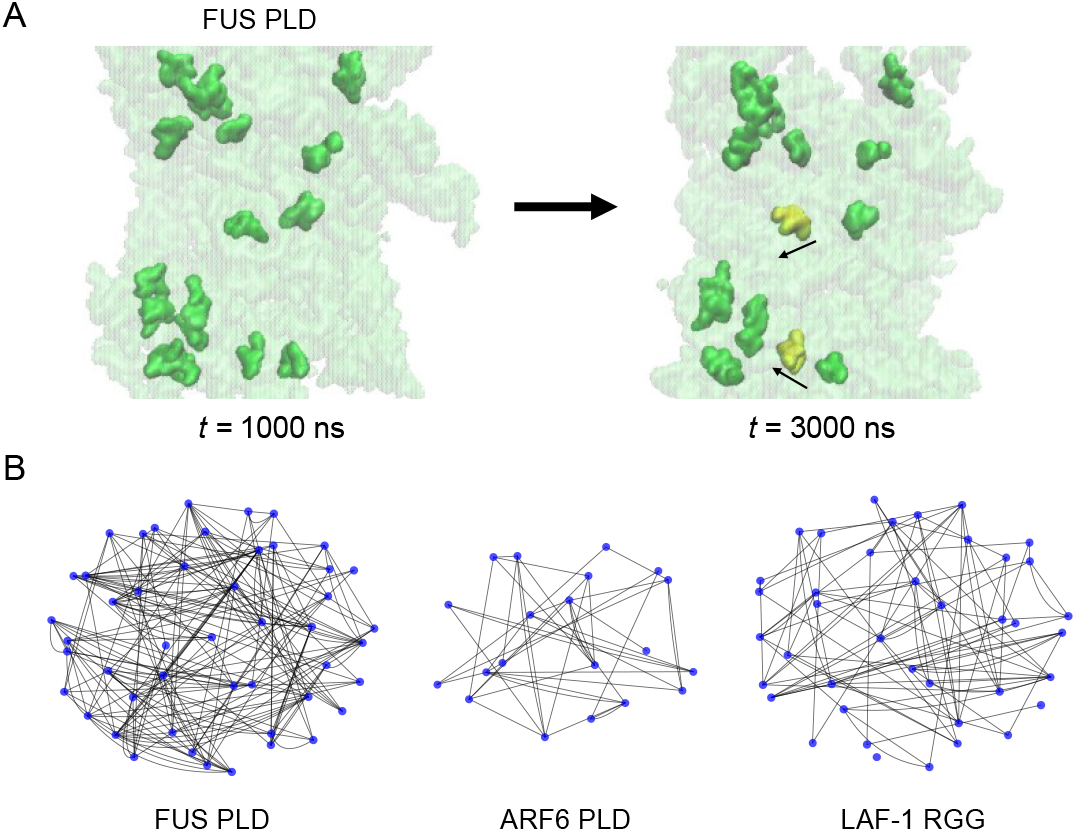
HOI-mediated *motif interaction*s within condensate and their multigraph representations. (A) Representative arrangement of HOI-mediated *motif interaction*s within FUS PLD condensate with 2 μs interval. About 1/4 *motif interaction*s are shown explicitly. Only a small fraction of *motif interaction*s (colored in yellow) undergoes slight slippage (black arrows). (B) Multigraph representations for FUS PLD (left panel), ARF6 PLD (middle panel) and LAF-1 RGG (right panel) condensates. Nodes and edges represent individual IDPs and HOI-mediated *motif interaction*, respectively. The number of edges between a pair of nodes can be more than 1, corresponding to multiple HOI-mediated *motif interaction*s between a IDP pair.

The schemes of the HOI-crosslinked network are shown in Fig. 2B. Nodes and edges represent individual IDPs located in the bulk or interface of condensate and HOI-mediated *motif interaction*, respectively. The number of edges between a pair of nodes can be more than 1, corresponding to the presence of multiple HOI-mediated *motif interaction*s between a IDP pair. Accordingly, such HOI-crosslinked network can be characterized as a multigraph (70).

The degree of nodes (the number of HOI-mediated *motif interaction* of the given IDP with the other IDPs) is 8.23 (FUS PLD), 4.19 (ARF6 PLD) and 7.14 (LAF-1 RGG). It is much lower than the residue contacts of a IDP (284.2, 145.2, 226.1, respectively), while these sparse HOI-mediated *motif interaction*s dominate the interaction strength of IDP. Besides, the degree of nodes representing IDP in condensate interface is lower than nodes in the bulk: 7.19 vs. 8.73 (FUS PLD) and 6.11 vs. 7.34 (LAF-1 RGG). The difference reflects the change of the number of HOI-mediated *motif interaction*s when one IDP moves from the condensate bulk to the interface (12). It should be noted that the network topology is quite stable during the μs-simulation (Fig. 2A and Fig. S10). That corresponds to the steady arrangement of HOI-mediated *motif interaction* within condensates.

### Estimation of condensate viscoelasticity based on our model

The reconfiguration dynamics of the HOI-crosslinked network was analyzed. And the network reconfiguration time (τ_r_) is dependent on the number of HOI-mediated *motif interaction* in condensates (*n*) and their lifetimes (τ_m_) (SI Ap*pendix*). Both can be directly obtained from simulations (Fig. 3A and *SI Appendix*). The corresponding expression is given by

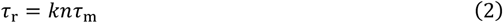

where *k* is determined by the protein length and concentration (*SI Appendix*). The estimated τ_r_ of three condensates are 0.12 ms (FUS PLD), 0.03 ms (ARF6 PLD) and 0.29 ms (LAF-1 RGG) (Fig. 3A). Notably, the reported relaxation time of the viscoelastic moduli of condensates is typically on the millisecond timescale (33, 34). For example, the relaxation time of the condensate formed by the low-complexity domain of the RNA-binding protein hnRNP A1 (A1-LCD) is ∼1 ms (33). The consistency with the experimental result suggests the inherent relaxation dynamics of condensate can be depicted as the reconfiguration of HOI-crosslinked network (34, 71, 72).

**Fig. 3.**
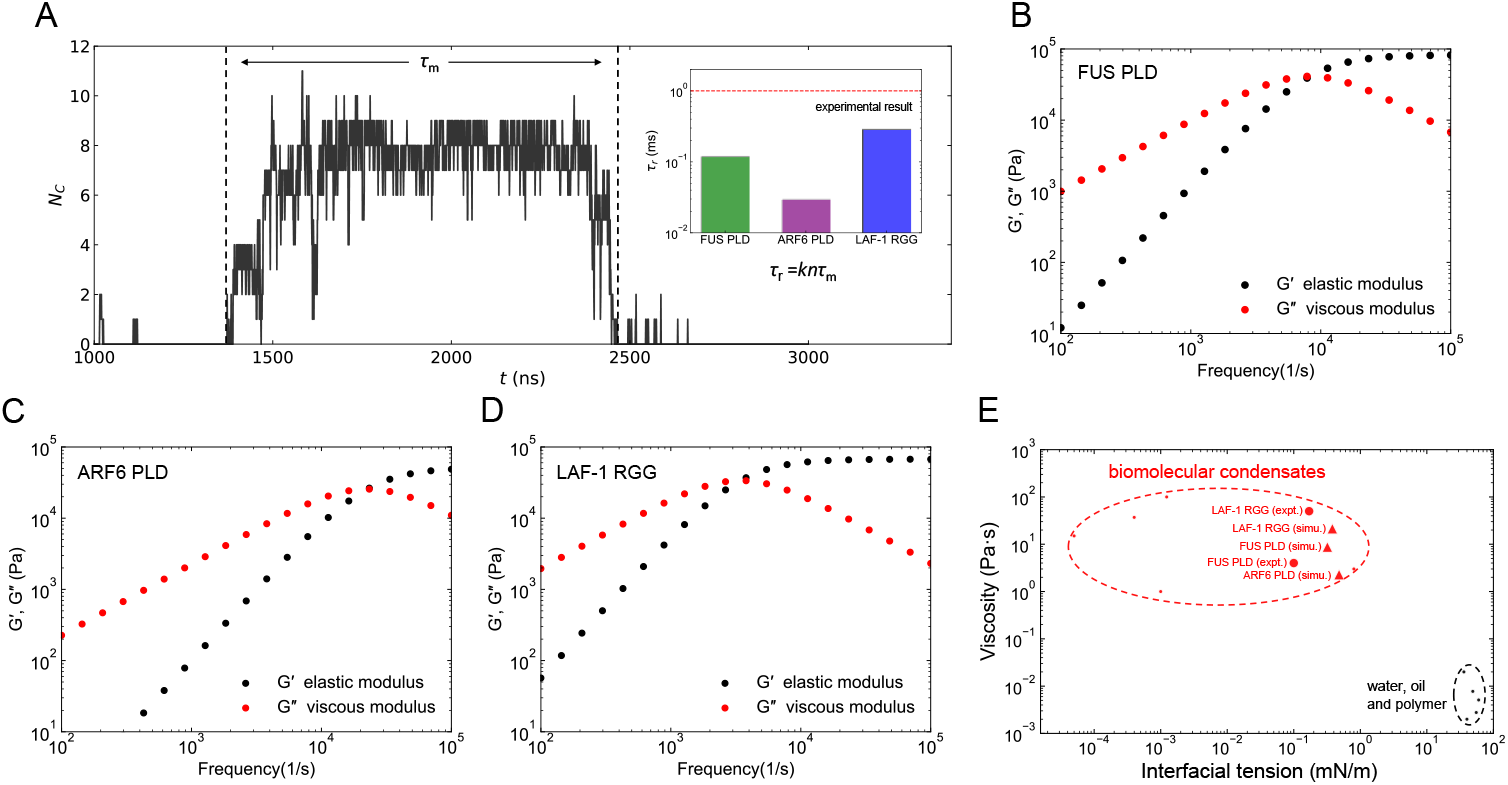
Estimation of condensate viscoelasticity and interfacial tension based on our model. (A) Representative time revolution of the number of residue contacts between HOI-mediated *motif interaction*. This *motif interaction* makes at *t* ≈ 1 50 ns and breaks at *t* ≈ 2450 ns. And the corresponding lifetime (τ_*m*_) is about 1100 ns. Estimated relaxation times (τ_*r*_) of three condensates are shown in the inset. The red dashed line indicates experimental results of relaxation times (33, 34). (B, C and D) Estimated viscoelastic moduli of three condensates: FUS PLD (C), ARF6 PLD (D) and LAF-1 RGG (E). (E) Estimated viscosities and interfacial tension coefficients of three condensates. The estimated values are shown in red triangles and the experimental results are shown in red solid circles.

The frequency dependence of the viscoelastic moduli (*G*(ω) = *G*^’^ + *iG*^′′^) was then estimated based on our network model (*SI Appendix*). The storage modulus (*G*^′^) characterizes the elasticity of condensate, and the loss modulus (*G*^′′^) characterizes the viscosity of condensate (30, 34). The viscoelastic modulus is given by (30, 73)

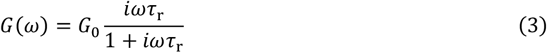

where *G*_0_ is the characteristic elastic modulus of condensate determined from temperature, protein length and concentration (*SI Appendix*). The viscoelastic moduli of three condensates are shown in Fig. 3B, C and D. At *ω* < 1/τ_r_, *G*^′′^ is larger than *G*^′^, indicating that the viscoelastic property of the condensates is predominantly viscous. And at *ω* > 1/τ_r_, *G*^′^ surpasses *G*^′′^, indicating predominantly solid-like behavior. The frequency of crossover between predominantly viscous and predominantly elastic is *ω*_c_ = 1/τ_r_ and the corresponding viscoelastic modulus is *G*_c_.

Based on the estimated viscoelasticity modulus at crossover frequency *G*_*c*_, the viscosity η of condensate can be obtained (30, 34),

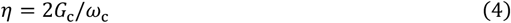

The predicted viscosities for three condensates are 8.8 Pa·s (FUS PLD), 2.3 Pa·s (ARF6 PLD) and 21.5 Pa·s (LAF-1 RGG) (Fig. 3E). These values fall within the reported range of viscosity of condensates (≥0.1 Pa·s, Fig. S1). Furthermore, the estimated viscosities for both FUS PLD and LAF-1 RGG condensates are consistent with their experimental measurements (FUS PLD: ∼4 Pa·s (46, 74); LAF-1 RGG: 10∼50 Pa·s (53, 75)).

Taken together, we propose that the protein condensate can be conceived as a physically crosslinked network mediated by higher-order residue interactions between finite motifs. And the steady arrangement of HOI-mediated *motif interaction*s with ultralong lifetime can result in the resilient configuration of condensate. Our model provides a methodology to calculate the viscoelastic moduli of condensate without any adjustable parameters and directly reproduce experimental results of relaxation time and viscosity.

### Estimation of condensate interfacial tension based on our model

As mentioned above, HOIs between finite motifs dominate IDP interactions within condensate (*SI Appendix*, Table S4). Hence, the cohesive energy of IDP as well as the interfacial tension of condensate were estimated accordingly. We used the lifetime of *motif interaction* (τ_m_ = τ_a_ exp(ϵ) (64, 76)) to calculate its strength (ϵk_B_T), where τ_a_ represents the formation time of *motif interaction* (*SI Appendix*). Given the sparse arrangement of HOI-mediated *motif interaction*s (Fig. 1F), the cohesive energy of IDP is weak. Therefore, the energy penalty for IDPs moving from condensate bulk to the interface is small and should result in very low interfacial tension f condensate. The interfacial tension coefficient (γ) can be expressed by (*SI Appendix*)

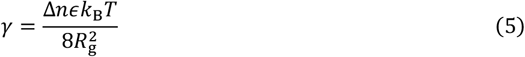

where *R*g is the radius of gyration of IDP and Δ*n* is the difference in the number of HOI-mediated *motif interaction*s between condensate bulk and the interface (*SI Appendix*). The estimated γs of three condensates are 0.32 mN/m (FUS PLD), 0.48 mN/m (ARF6 PLD) and 0.38 mN/m (LAF-1 RGG) (Fig. 3E). These values fall within the reported range of interfacial tension coefficient of condensates (<1 mN/m, Fig. S1). Furthermore, the estimated interfacial tension coefficients for both FUS PLD and LAF-1 RGG condensates are consistent with their experimental measurements (FUS PLD: ∼0.1 mN/m (31); LAF-1 RGG: 0.17 mN/m (75)). In summary, the proposed HOI-crosslinked network can also reproduce experimental results of interfacial tension coefficient of condensate.

### Cooperative residue interactions within *motif interaction*

The details of higher-order residue interactions within three condensates were studied. Fig. 4A shows representative configurations of the interaction between residue motifs 151-153 and 157-160 in FUS PLD condensate. They include 4 glutamines (Q): the residues can interact with multiple residues, and their multivalent interactions undergo local reorganization. For example, at *t* = 1444.7 ns, Q153 forms hydrogen bonds with Q158 and Q160, and Q160 forms hydrogen bonds with P152 and Q153. At *t* = 1445.6 ns, the hydrogen bond between Q153 and Q160 is substituted by hydrogen bond between P152 and Q160. Meanwhile, the hydrogen bond between Q153 and Q158 remains.

**Fig. 4.**
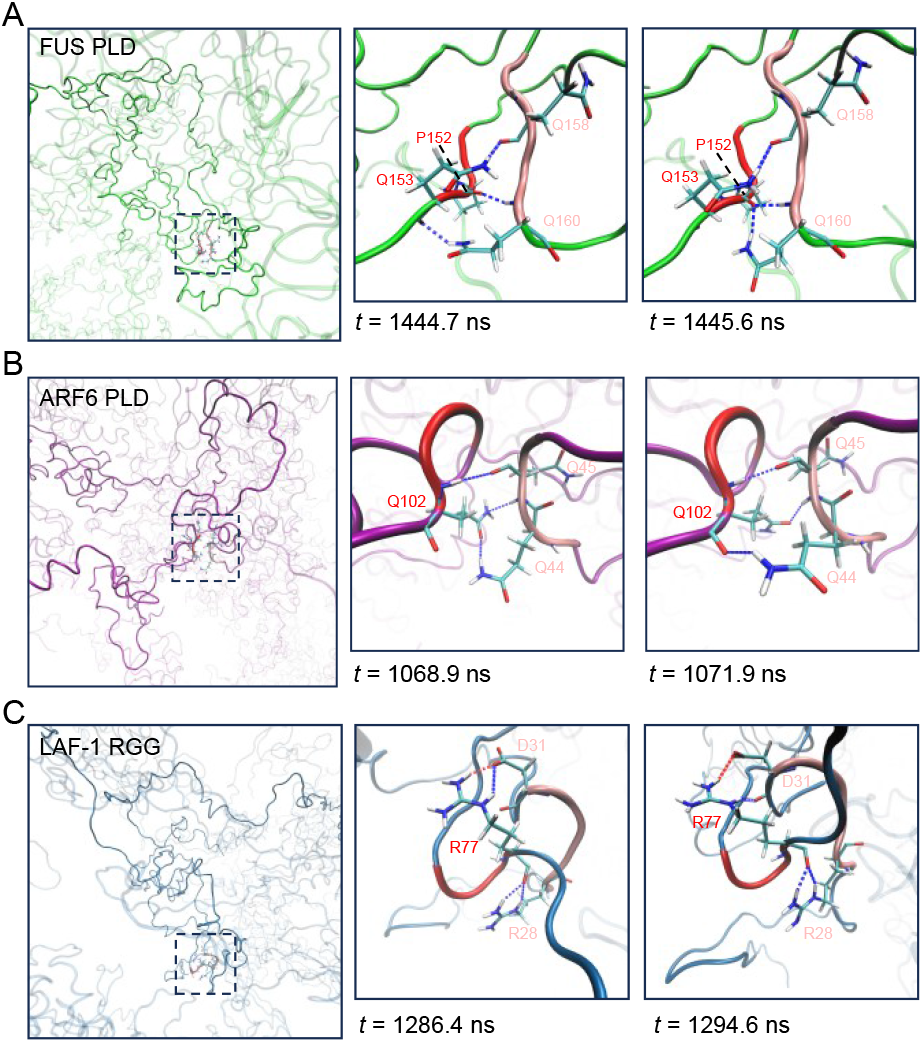
Higher-order residue interaction between HOI-mediated *motif interaction* in three condensates. (A, B and C) Representative configurations of multivalent residue interactions and their localized reorganization within FUS PLD (A), ARF6 PLD (B) and LAF-1 RGG (C) condensate. Hydrogen bonds and salt bridges are shown in blue and red dashed lines, respectively.

We further identified other type of higher-order residue interaction within FUS PLD condensate (Fig. S11), e.g., the case including 2 tyrosine and 1 glutamine (141-143 and 4-6). Similarly, the residues have multivalent interaction and the remaking of residue interaction is localized. On average, the number of hydrogen bonds between residue motifs (with 3.79 residues) is 2.54.

Fig. 4B and C display representative snapshots of HOI-mediated *motif interaction*s within ARF6 PLD and LAF-1 RGG condensates, respectively. Additional higher-order residue interactions were identified, e.g., the cases including 6 glutamine (100-102 and 44-46, ARF6 PLD), 2 arginine and 2 aspartic acid (75-77 and 28-31, LAF-1 RGG). The residue compositions of these HOIs are quite diverse. Interestingly, they can similarly form sufficient interaction which engenders the localization of interaction remaking. For ARF6 PLD condensate, there are 2.59 hydrogen bonds between the motifs with 3.57 residues. As for LAF-1 RGG condensate, the numbers of hydrogen bonds and salt bridges in HOI-mediated *motif interaction*s are 2.15 and 0.53, respectively. And the average length of residue motifs is 4.09.

Overall, higher-order residue interactions are prevalent in IDP condensates which result in sufficient residue contacts and localized reorganization within finite residue motifs. Therefore, the corresponding residue motifs can maintain ultralong-lasting interaction. In FUS PLD condensate, tyrosine and glutamine can form multivalent residue interactions and are important to higher order interaction. And HOIs within ARF6 PLD condensate are mainly attributed to glutamine. As for LAF-1 RGG condensate, HOIs are attributed to arginine, tyrosine and aspartic acid (Fig. S12).

Combinations of these important residues in HOI-mediated *motif interaction* and their probabilities are summarized in Table S6. In FUS PLD condensate, stoichiometric ratios of the top four most populated combinations are 1:2 (Y:Q, the same below, 15.0%), 1:1 (13.3%), 2:1 (12.4%), 1:3 (8.3%). Notably, each residue motif tends to have multiple important residues to form higher-order residue interaction. As for ARF6 PLD condensate, the number of glutamines in the top four most populated combinations are 4 (25.6%), 2 (16.3%), 3 (16.3%), 6 (7%). In LAF-1 RGG condensate, stoichiometric ratios of the top four most populated combinations are 1:1 (R:D, 6.8%), 2:1:1 (Y:R:D, 5.6%), 2:1 (R:D, 5.1%), 1:1:1 (Y:R:D, 5.1%). For both condensates, most higher-order residue interactions tend to comprise multiple important residues from each residue motif. Accordingly, the consecutive sequence positions of the important residues can facilitate the cooperativity of their interactions and formation of higher order interaction (Fig. 1C, D and E; Fig. S9).

Sequence analysis of HOI-mediated *motif interaction* provides enriched information about the sequence features of IDP interaction (i.e., molecular grammar). In addition to well-conceived tyrosine and arginine (36-38), glutamine and aspartic acid also play crucial role in IDP interaction (Table S6; Fig. S12). Our findings highlight the importance of sequence position of these residues. Systematic studies about sequence features of higher-order residue interactions should open a new avenue for deep deciphering the molecular grammar of IDP interaction.

## Discussion

We systematically studied the IDP interaction within condensates based on μs-scale atomistic molecular dynamics simulations of three homogeneous condensate (FUS PLD, ARF6 PLD and LAF-1 RGG condensate), as well as the reported atomistic simulation trajectories of two heterogeneous condensate (H1-ProTα condensate (55) and protamine-ProTα condensate (23)). As revealed by our results, multivalent residue interactions emerge in the finite residue motifs between all interacting IDP pairs of these condensates. In addition, the reorganization of residue interaction is localized and their interactions are apparently non-additive. Such higher-order residue interactions contribute to the extremely long lifetime (on the order of μs timescale) of the interaction between corresponding residue motifs which is denoted as HOI-mediated *motif interaction*.

Notably, the interaction energy between IDPs is dominated by HOI-mediated *motif interaction*s. Versatile types of cooperative residue interactions were identified. In FUS PLD and LAF-1 RGG condensate, tyrosine and arginine can lead to cooperativity based on π-interaction (Fig.S11), which is similar to the case of sticker-and-spacer model (36-38). In addition, networks of hydrogen bonds involving multiple glutamines were also observed in FUS PLD and ARF6 PLD condensate. Cooperativity can be also attributed to the hybrid interactions of hydrogen bond and salt bridge mediated by arginine and aspartic acid were identified in LAF-1 RGG condensate. Taken together, our results reveal a spectrum of cooperative residue interactions and their sequence features. In addition to tyrosine and arginine, our findings illustrate the importance of glutamine and aspartic acid. Besides, the consecutive sequence position of these important residues is also essential for these highly cooperative interactions. Interestingly, the impacts of sequence spacing of important residues on IDP interactions have been reported in previous study (37): shortening the spacing of tyrosine within A1-LCD sequence can facilitate IDP interaction (37). Our findings about sequence features for higher-order interaction can enrich the insight into the molecular grammar of IDP interaction.

HOI-mediated *motif interaction*s and the cohesive energy of IDP can be also used to interpret the saturation concentration of IDP (*c*_sat_) via (77),

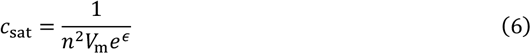

where *V*_m_ is the volume of *motif interaction* (*SI Appendix*). The estimated saturation concentrations of FUS PLD and LAF-1 RGG are 114 μM and 53 μM, respectively, which are consistent with their experimental results (FUS PLD: ∼125 μM (36); LAF-1 RGG: 12 μM (78)).

Notably, the spatial arrangement of HOI-mediated *motif interaction*s within condensate is quite stable. IDP condensate can be conceived as a HOI-crosslinked network. The viscoelastic moduli along with the relaxation time and viscosity can be calculated and the estimated values are consistent with experimental results. The steady arrangement of HOI-mediated *motif interaction* results in slow reconfiguration of condensate, which leads to ultrahigh viscoelasticity. On the other hand, the sparse arrangement of HOI-mediated *motif interaction* leads to the weak cohesive energy of IDP, resulting in very low interfacial tension of condensate. The estimations of interfacial tension coefficients, along with saturation concentration, are also consistent with the experiment. In summary, our proposed HOI-crosslinked network successfully reproduces the experimental results of condensate viscoelasticity, interfacial tension and saturation concentration, providing a unified and quantitative interpretation of seemingly diversified physical properties of condensate.

## Materials and Methods

FUS PLD, ARF6 PLD and LAF-1 RGG refer to the low-complexity prion-like domain of Fused in Sarcoma (1-165 (79-81)), the prion-like domain of auxin response factor 6 (464-618 (82, 83)) and R/G-rich domain of LAF-1 (1-168 (5, 60)), respectively. Atomistic MD simulations were performed using GROMACS (84) 2022.6 or newer. See *SI Appendix* for detailed description of simulation protocol and analysis methods.

## Acknowledgments

This work was supported by the National Natural Science Foundation of China (12175195 and 32371299). We acknowledge Haiping Fang, Xiandeng Wu and Hai Lei for valuable feedback. We thank Shi-Jie Chen, Changsong Zhou, Ruoyao Zhang, Haoyu Song and Xialan Zhang for helpful discussions. We also thank Benjamin Schuler, Robert B. Best and Wenwei Zheng for atomistic simulation trajectories. Major computations were done using Quantum Many-Body Simulation HPC.

## Notes

### Competing Interest Statement

The authors have declared no competing interest.

