## Supplementary material for "Higher-order Residue Interactions Encode Diversified Physical Properties of Biomolecular Condensates": Figure. S1

**This PDF file includes:**

Supporting text

Figures S1 to S12

Tables S1 to S6

SI References

Supporting Information Text

**Proteins and corresponding sequences used in this work**

Throughout this paper, FUS PLD, ARF6 PLD and LAF-1 RGG refer to the low-complexity prion-like domain of Fused in Sarcoma (1-165 (1, 2)), the prion-like domain of auxin response factor 6 (464-618 (3, 4)) and arginine/glycine-rich domain of LAF-1 (1-168 (5, 6)), respectively. FUS PLD and LAF-1 RGG are commonly used as model systems for investigating the mechanism underlying biomolecular condensation and related physical properties (6-9). ARF6 belongs to the ARF family of transcription factors, which can modulate auxin responses to coordinate normal plant growth and development (3, 4). The PLDs in ARF protein have been shown to be necessary for condensate formation and associated cellular processes (3, 4). The sequences of these three IDPs are present below.

FUS PLD

MASNDYTQQATQSYGAYPTQPGQGYSQQSSQPYGQQSYSGYSQSTDTSGYGQSSYSSYGQSQNTGYGTQSTPQGYGSTGGYGSSQSSQSSYGQQSSYPGYGQQPAPSSTSGSYGSSSQSSSYGQPQSGSYSQQPSYGGQQQSYGQQQSYNPPQGYGQQNQYNSSS

ARF6 PLD

QFQNSPGFSMQSPSLVQPQMLQQQLSQQQQQLSQQQQQQQQLSQQQQQQLSQQQQQQLSQQQQQQLSQQQQQQAYLGVPETHQPQSQAQSQSNNHLSQQQQQVVDNHNPSASSAAVVSAMSQFGSASQPNTSPLQSMTSLCHQQSFSDTNGGNNP

LAF-1 RGG

MESNQSNNGGSGNAALNRGGRYVPPHLRGGDGGAAAAASAGGDDRRGGAGGGGYRRGGGNSGGGGGGGYDRGYNDNRDDRDNRGGSGGYGRDRNYEDRGYNGGGGGGGNRGYNNNRGGGGGGYNRQDRGDGGSSNFSRGGYNNRDEGSDNRGSGRSYNNDRRDNGGDG

**molecular dynamics (MD) simulations**

Atomistic simulations of condensate systems formed by FUS PLD, ARF6 PLD and LAF-1 RGG were conducted using GROMACS (10), version 2022.6 or newer. For the atomistic simulations, the Amber99SB-disp force field with the modified TIP4P-D water model was adopted (11). The combination of Amber99SB-disp protein force field and the accompanying water model can reproduce well the available experimental results of IDP conformational ensembles (11-14) and have been wildly utilized for the atomistic simulations of IDP condensates (15-17). The temperature was kept constant at 300 K using velocity-rescaled thermostat (18), and the pressure was kept at 1 atm with the Parrinello-Rahman barostat (19). The periodic boundary conditions were adopted in all directions (20). Both Van der Waals interactions and short-range repulsion were calculated with a cutoff at 1.2 nm (11). Beyond this distance, the long-range electrostatic interactions were modelled using the particle mech Ewald method (21). Covalent bonds involving hydrogen atoms were constrained through the LINCS algorithm and equations of motion were integrated under a time step of 2 fs (22).

The initial configurations for atomistic simulations of FUS PLD condensate in slab geometry were generated from a coarse-grained (CG) simulation on the basis of the procedure described previously (23-25). The Martini3-IDP model was used for CG simulation, which can reproduce condensate formation via phase separation for both homotypic and heterotypic IDP-based systems (26). The CG simulations was conducted in accordance with the standard protocol (26). At first, 48 chains of FUS PLDs were randomly inserted in a box of size 14×14×35 nm^3^ to serve as the Initial dispersed configuration. The simulation box was then solvated by water molecules with the salt concentration of 150 mM. Subsequently, the CG system was subjected to 50,000 steps of energy minimization, and successively equilibrated under the NVT (300 K) ensemble and the NPT (300 K and 1 atm) ensemble both for 200 ns. The production of CG simulation was conducted in the NVT ensemble at 300 K for a duration of 10 microseconds to generate a phase-separated system. Three independent CG simulations for FUS PLD condensate were performed and each final configuration (proteins only) was mapped to atomistic structure with the Backward algorithm (26). This algorithm includes the elimination of side-chain clashes in the atomistic representation (26).

For the atomistic simulation of ARF6 PLD condensate in cubic geometry, the initial configuration was also generated from CG simulation using the same procedure mentioned above. The CG system comprises 20 chains of ARF6 PLD, with a box size of 13×13×13 nm^3^ (~15 mM) and a 5-μs NVT for producing phase-separated system was carried out. Besides, the initial configuration of atomistic simulation of LAF-1 RGG condensate was taken from a previously reported atomistic simulation (23) (i.e., the final configuration of the trajectory) and the box size was set to 10×10×34 nm^3^. The atomistic configurations were also obtained using Backward algorithm. In the next step, the simulation box was solvated by TIP4P-D water molecules (11), for each atomistic system (i.e., three independent FUS PLD systems, one ARF6 PLD system and one LAF-1 RGG system). Sodium and chloride ions were then inserted to neutralize the solvated system and mimic the physiological conditions (150 mM NaCl). Each system was then energy-minimized and equilibrated successively with the NVT ensemble (300 K) and the NPT ensemble (300K, 1atm) both for 50 ns. Finally, a 3-μs production run in the NPT ensemble was carried out for each system. The first 1 μs was treated as system equilibration and was removed prior to analysis. The setup details of all condensate system are summarized in Table S2.

**Analysis of MD simulations**

Density profiles and residue pairwise contacts were both calculated with codes written in Python-3.10.9 using the trajectory analysis package MDAnalysis-2.9.0 (27, 28). The condensate interface is defined based on the mass concentration of proteins (29). A residue contact is defined when the minimum distance between any two heavy atoms of two residue is less than 0.4 nm. An interacting IDP pair forms when the number of residue contacts between them is over 30. All snapshots were drawn and rendered using VMD-1.9.3 (30).

The power spectral density (PSD) of the evolution of the number of residue contacts (*N*(*t*)) is given by (31)

$$\begin{aligned} \mathrm{PSD}=\lim_{T\to\infty} \frac{1}{T}\left\langle\left| \int_{O}^{T} N_{c}\left( t \right)e^{iwt}dt \right|^{2} \right\rangle\#\left( 1 \right) \end{aligned}$$

where the angle brackets refer to the ensemble average. PSD is related to the autocorrelation function (i.e., $<N(t)N(t+\tau)>$).

The example of making and breaking of *motif interaction* is shown in Fig. 3A. As *motif interaction* forms, residue contacts begin to occur between corresponding residue motifs. Accordingly, the formation time of *motif interaction* $\tau_{a}$ can be approximately equated to the lifetime of normal residue contacts (~1 ns). When *motif interaction* breaks, no residue contacts between two motifs remain. The lifetime of *motif interaction* was calculated based on the entire trajectory (i.e., 3000 ns). *Motif interaction*s are considered to break at *t* = 3000 ns, if they remain until the end of the simulation.

The interaction energies between IDPs were calculated by postprocessing the MD trajectories with the *mdrun -rerun* routine in GROMACS (10). The distance cutoffs for coulomb and van der Waals interactions were set to 3.5 nm (7). The interaction energies associated with *motif interaction*s include all energy contributions involving the corresponding residue motifs (Table S4).

The calculation of hydrogen bond and salt bridge was based on the geometrical configuration. Hydrogen bond is defined if the distance between donor (D) of H-bond and acceptor (A) is less than 0.35 nm and the angle of H-D···A is lower than 40°. Salt bridge is defined if the distance between the nitrogen atom of positively charged residues and oxygen atom of negatively charged residues is less than 0.4 nm. They were both calculated using the package MDAnalysis (27, 28).

**Construction of HOI-crosslinked network and calculations for physical properties**

*Construction of HOI-crosslinked network.* HOI-crosslinked network was constructed on the basis of the theory of associative polymers (32, 33). The idea of this theory has been used to describe the driving force for phase separation of PLDs, i.e., sticks-and-spacers model (34-36). The theory we deployed in this paper considers a solution of multivalent polymers, with associating groups which can form sustained HOI-mediated *motif interaction*s. The size of each associating group (i.e., monomer) is identical to that of an individual residue motif unit with the length of ~4 (i.e., 4-residue segment). The lifetime and interaction energy of associating groups are $\tau_{m}$ and $\epsilon k_{B}T$, where $\epsilon$ refers to a scaling parameter, $k_{B}$ is the Boltzmann’s constant and $T$ is temperature. The relation of $\tau_{m}$ and $\epsilon$ is $\tau_{m}=\tau_{a}\exp\left( \epsilon\right)$ (37, 38), where $\tau_{a}$ represents the formation time of *motif interaction*. Each chain has $n$ residue motifs separated by spacers of $l\approx N/4n$ monomers each, where $N$ is the number of residues in IDP. The concentration of 4-residue segments is $c$, and the monomer volume fraction is $\phi=cV_{m}$, where $V_{m}$ is the volume of *motif interaction*. The monomer volume fraction $\phi$ should be larger than the gelation point $\phi_{g}$, because IDPs usually undergo phase separation coupled to percolation (32, 39-42).

*Calculations for viscoelastic moduli and viscosity*. According to our finding, the condensates we considered can be conceived as a physically crosslinked network mediated by HOI-mediated *motif interaction* between IDPs. The reconfiguration of the network can be considered to be dominated by making and breaking of *motif interaction* (33, 43, 44). And relation between network reconfiguration time and lifetime of *motif interaction* is given by (33),

$$\begin{aligned} \tau_{r}\approx\tau_{m}n^{2}l^{2+2z}\phi^{\frac{2+2z}{3v-1}}\#\left( 2 \right) \end{aligned}$$

where *z* and *v* are exponents related to the types of solvent. For θ-solvent, *z* = -0.5 and *v* = 0.5 (33, 45). And Eq. (2) can be written as,

$$\begin{aligned} \tau_{r}\approx\frac{N}{4}\phi^{2}n\tau_{m}=kn\tau_{m}\#\left( 3 \right) \end{aligned}$$

where $k = \frac{N}{4}\phi^{2}$ and is determined by the chain length and concentration.

There are several typical relaxation times which should influence the dynamic moduli of system (33), including the diffusion time of monomer (46), the single-chain relaxation time (46-48), and the reconfiguration time of the whole system (43, 44, 49). And the viscoelastic properties of ideal polymer solutions should conform to a generalized Maxwell model of all these relaxation times (48, 50, 51). Notably, the reconfiguration time (≥ ms (44, 51)) is considerably larger than other relaxation times (≤ μs (24, 47)), and we focus on the condensate viscoelasticity on the time scale of reconfiguration time. According to the Maxwell model, the frequency dependence of the viscoelastic moduli is given by (50, 51)

$$\begin{aligned} G\left( \omega\right)=G_{0}\frac{i\omega\tau_{r}}{1+i\omega\tau_{r}}\#\left( 4 \right) \end{aligned}$$

where $G_{0}$ is the characteristic elastic modulus and $G_{0}=\frac{k_{B}T\phi}{NV_{m}}$ (33, 46). The frequency of crossover between predominantly viscous and predominantly elastic is $\omega_{c}=1/\tau_{r}$ and the corresponding viscoelastic modulus is $G_{c}$. Then the viscosity can be obtained (50, 51),

$$\begin{aligned} \eta=2G_{c}/\omega_{c} \#\left( 5 \right) \end{aligned}$$

*Calculations for interfacial tension*. Once phase separation occurs, a new surface with an associated interfacial tension created (52). Inside the condensate bulk, each chain shares a cohesive energy $u$, which is determined by the number and strength of HOI-mediated *motif interaction*s. Compared to the bulk, proteins at the condensate interface may loss some interactions and pay a penalty $\Delta$*u*. Accordingly, this penalty can be mainly attributed to the breaking of *motif interaction*s and thus $\Delta u=\Delta n\epsilon k_{B}T/2$. Here, $\Delta n$ refers to the difference in the number of *motif interaction*s between condensate bulk and the interface. For a protein of typical linear size *R*_g_, the energy penalty per unit area of the interface (i.e, interfacial tension) is given by (29),

$$\begin{aligned} \gamma\approx\frac{\Delta u}{\left( 2R_{g} \right)^{2}}=\frac{\Delta n\epsilon k_{B}T}{8R_{g}^{2}}\#\left( 6 \right) \end{aligned}$$

*Calculations for saturation concentration*. According to the theory of associative polymer (32, 36), the free energy of the system can be separated into two parts,

$$\begin{aligned} F=F_{\mathrm{HOI}}+F_{non-HOI}\#\left( 7 \right) \end{aligned}$$

where $F_{\mathrm{HOI}}$ and $F_{non-HOI}$ refer to the contributions of HOI-mediated *motif interaction* and other residues. Here, $F_{non-HOI}$ is directly determined by the solution condition, including chain length, concentration and interaction parameters between monomers. Thus, it can be seen as a constant for a fixed system. On the other hand, let $p$ be the faction of residue motifs that form HOI-mediated *motif interaction*, and $F_{\mathrm{HOI}}$ can be written as (32),

$$\begin{aligned} \frac{F_{\mathrm{HOI}}}{k_{B}T}=-\frac{cp}{2l}\ln\frac{c}{el}v_{b}+\frac{c}{2l}\left[ p\ln p+2\left( 1-p \right)\ln\left( 1-p \right) \right]-\frac{\epsilon cp}{2l}\#\left( 8 \right) \end{aligned}$$

Minimization of $F$ with respect to $p$ yields,

$$\begin{aligned} \frac{p}{\left( 1-p \right)^{2}}=\frac{c}{l}V_{m}e^{\epsilon}\#\left( 9 \right) \end{aligned}$$

Tree-like clusters without closed loops were assumed according to the Flory-Stockmayer theories (53, 54). Therefore, the gel point is defined when a single chain is connected to two neighboring chains. Through a further simplification, the absence of intramolecular interactions was assumed. These simplifications lead to $p\approx1/n$. Then we get the chain concentration at the gel point, the proxy for saturation concentration,

$\begin{aligned} c_{g}=c_{\mathrm{sat}}\approx\frac{1}{n^{2}V_{m}e^{\epsilon}}\#\left( 10 \right) \end{aligned}$

Figures

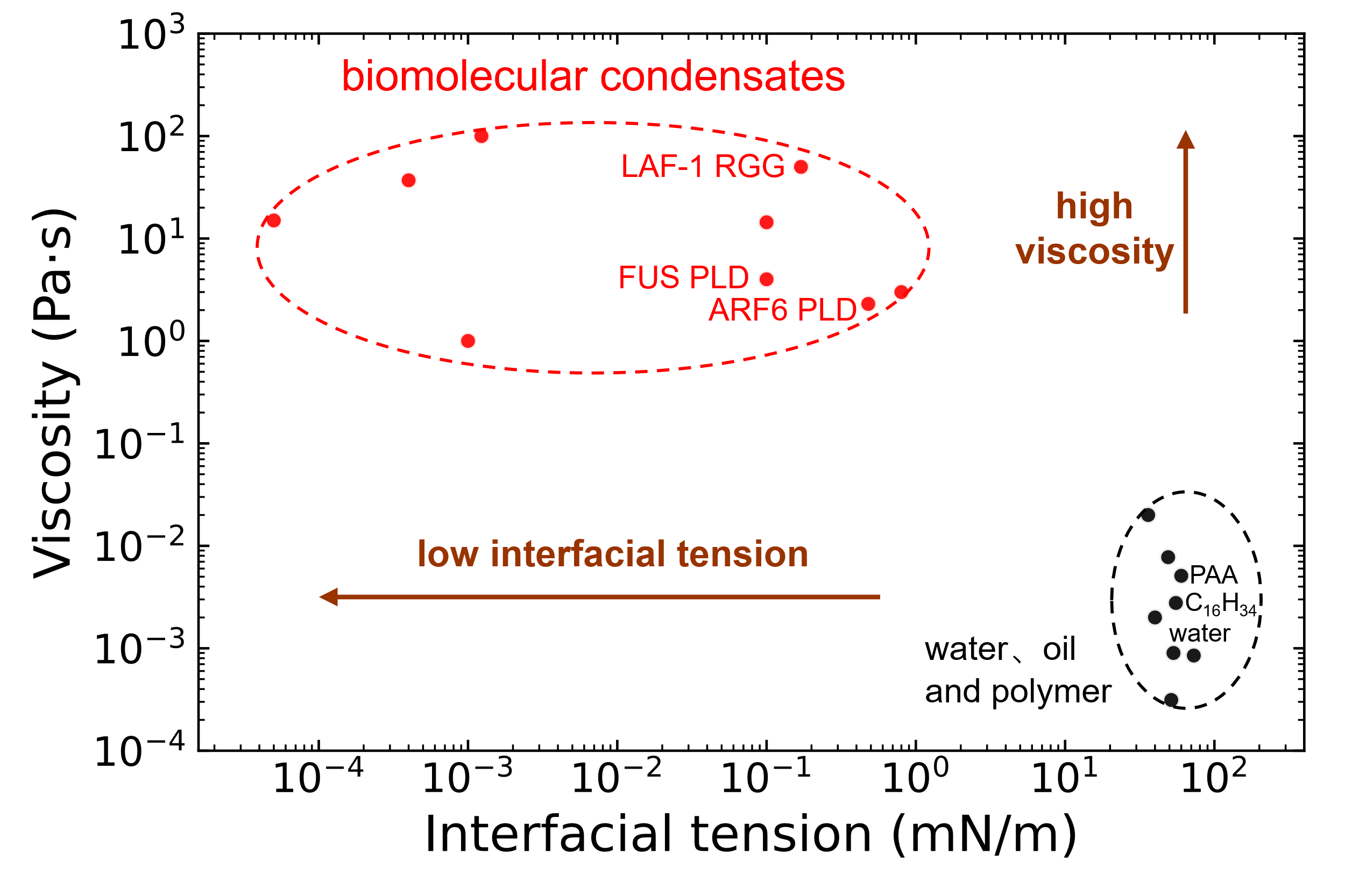
Figure. S1.

Fig. S1. Extraordinarily high viscoelasticity and low interfacial tension of biomolecular condensates. Compared with water, standard oil droplets and polymer, the viscosities of condensate are at least 100 times higher, whereas the interfacial tension coefficients are at least 100 times lower. See Table S1 for values and the corresponding references used in this plot.

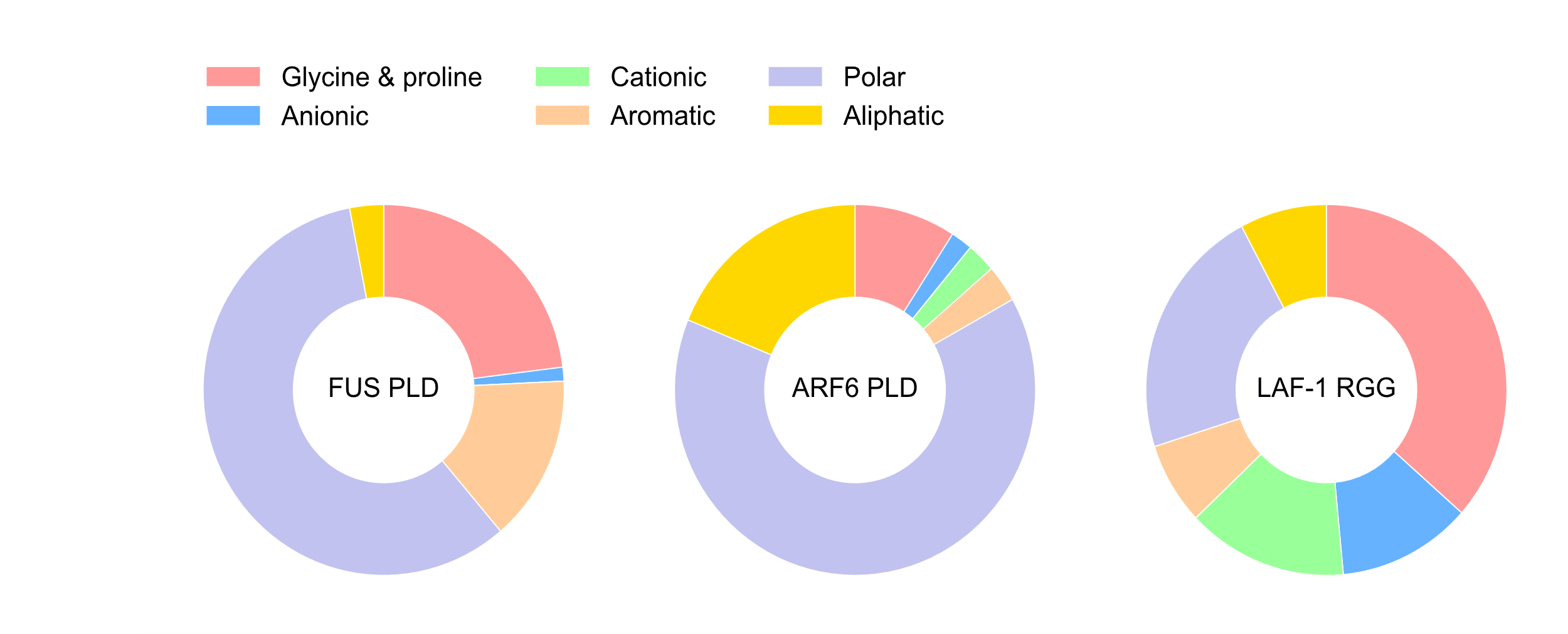
Figure. S2.

Fig. S2. Pie charts for the sequence composition of FUS PLD, ARF6 PLD and LAF-1 RGG. The composition is segregated into glycine and proline, cationic, polar, anionic, aromatic and aliphatic residues.

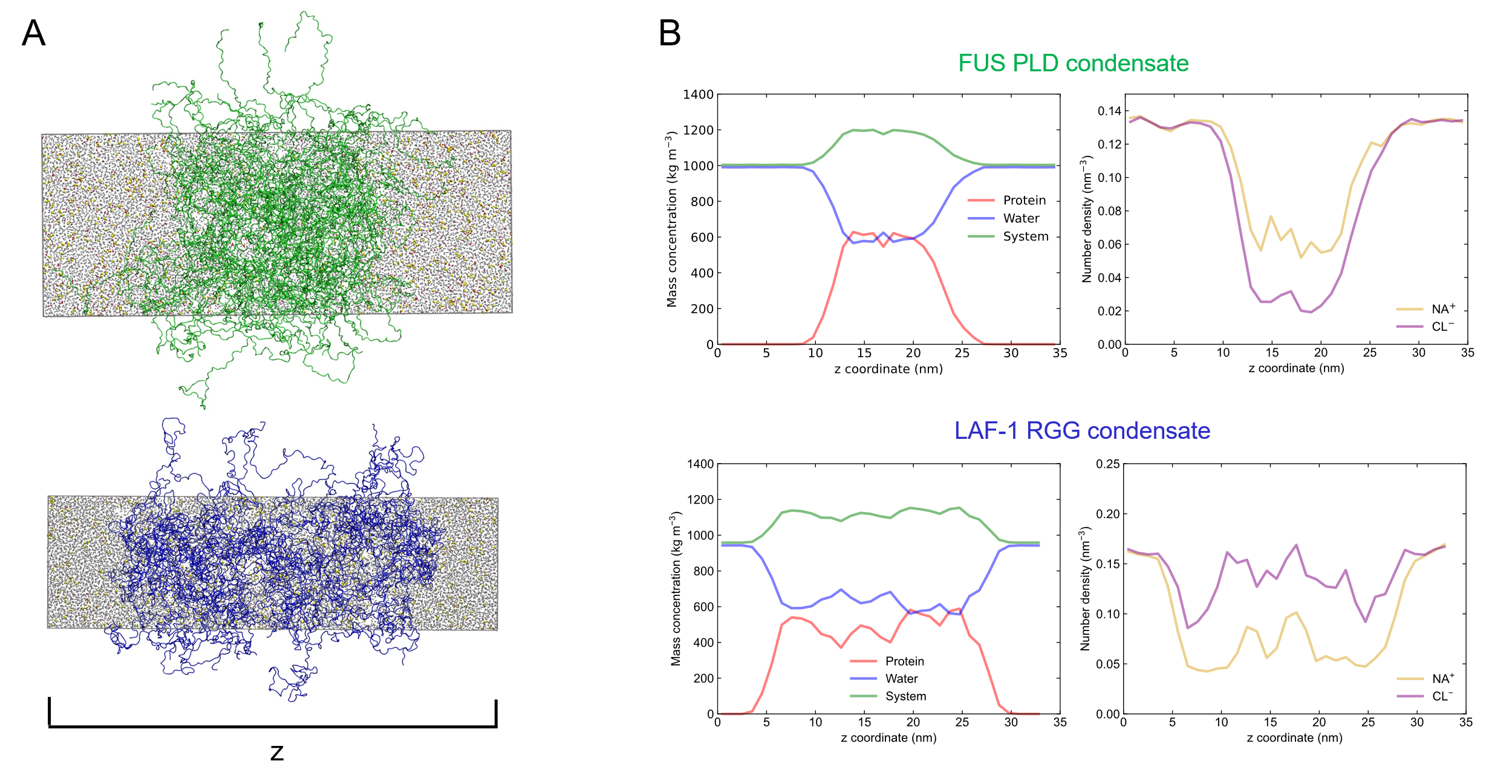
Figure. S3.

Fig. S3. Mass concentration and ion number density in MD simulations. (A) Representative snapshots of atomistic explicit-solvent simulation of FUS PLD (up) and LAF-1 RGG (bottom) condensates in slab geometries. Densities of all components were calculated along the z-axis of the simulation box. (B) Mass concentrations of protein, water and the system (including protein, water and ions; left panels), and number densities of Na^+^ and Cl^-^ ions (right panels). For FUS PLD condensate, the average density for three trajectories was taken.

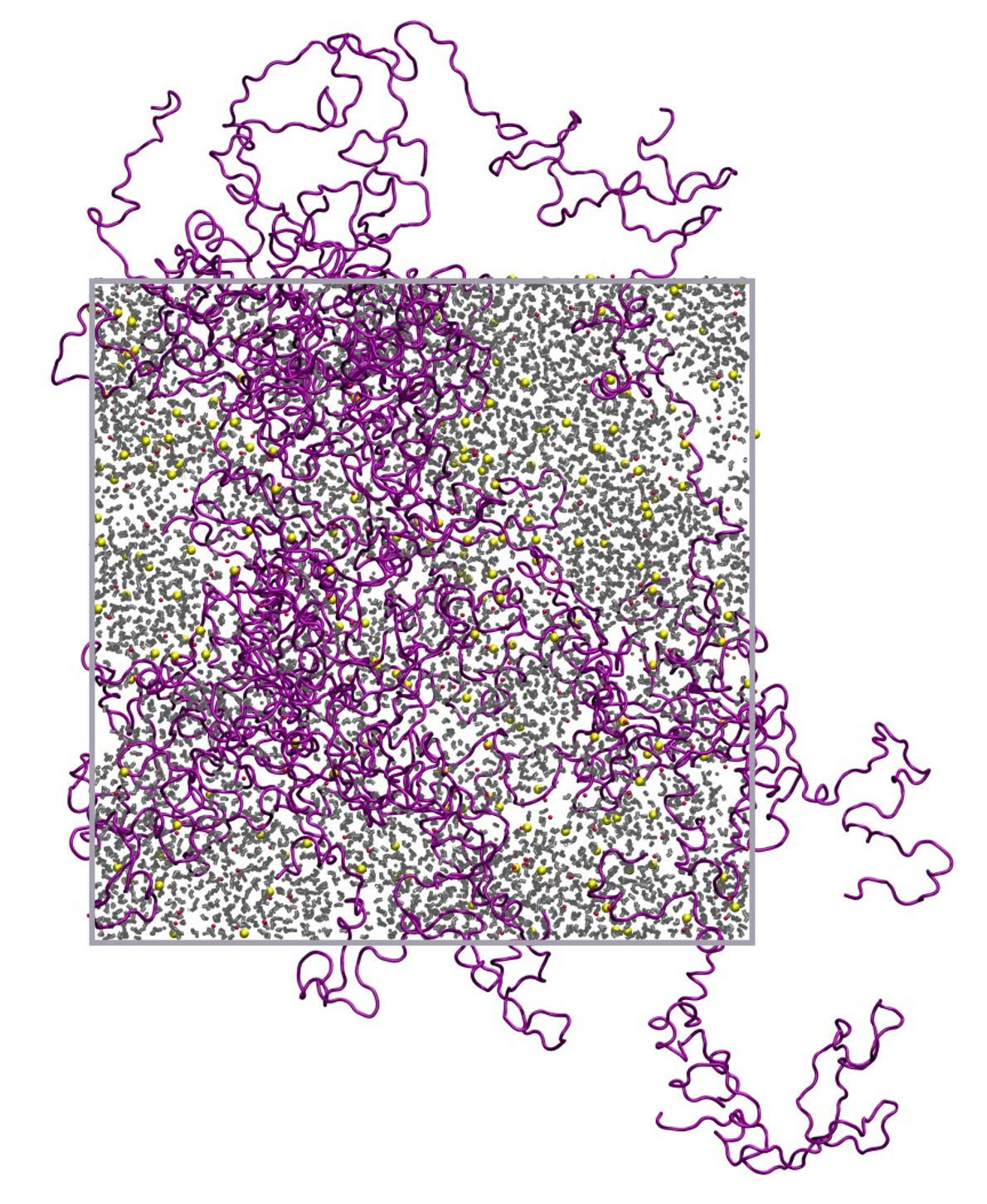
Figure. S4.

Fig. S4. Representative snapshot of atomistic explicit-solvent simulation of ARF6 PLD condensate in cubic geometry. Proteins are shown explicitly in purple strips, with water colored in gray, and Na^+^ and Cl^-^ ions in red and yellow, respectively.

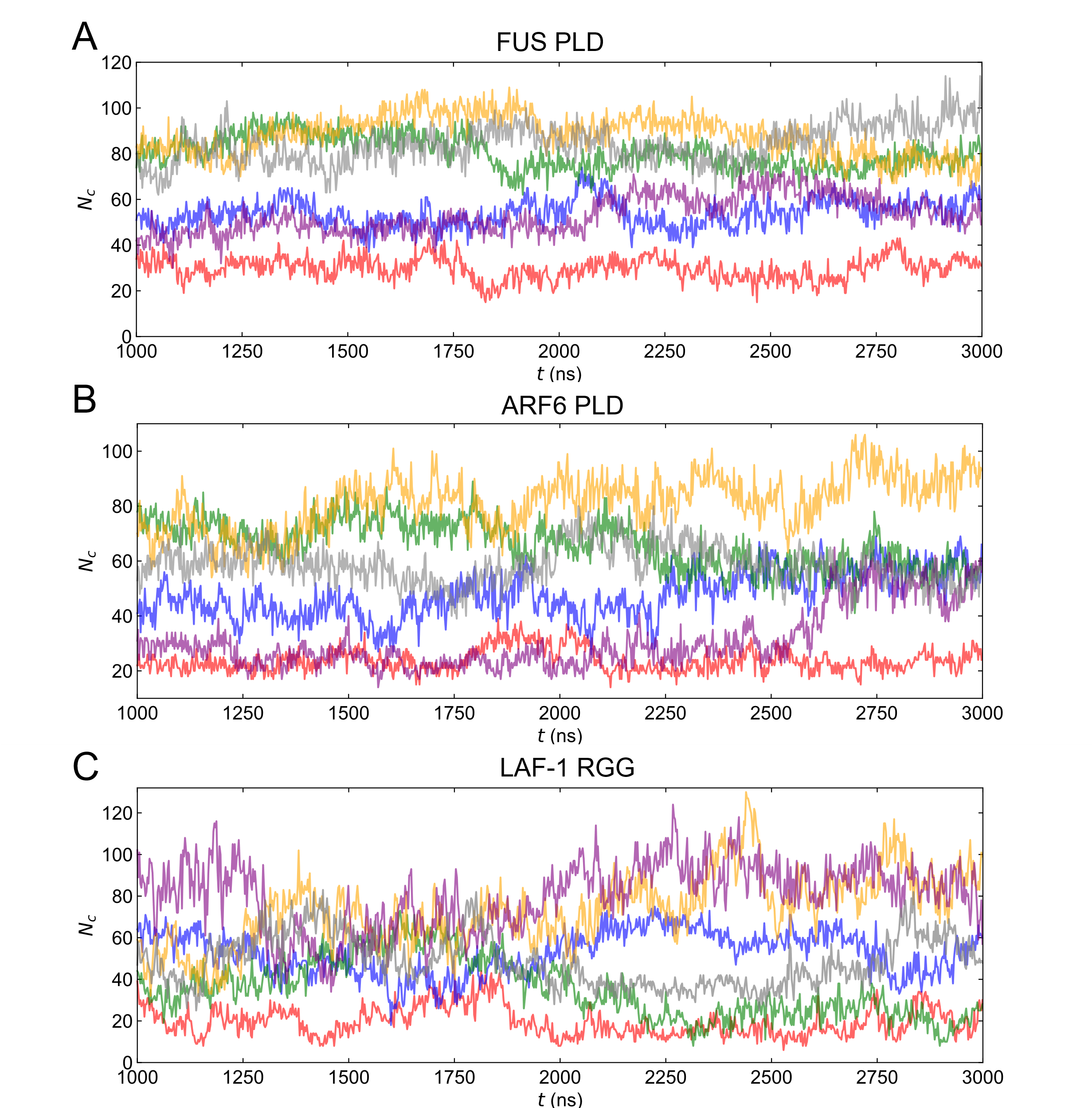
Figure. S5.

Fig. S5. The evolution of the number of residue contacts *N*_c_(*t*) between IDPs in FUS PLD, ARF6 PLD and LAF-1 RGG condensates. (A, B and C) Examples of contact number fluctuations between six interacting IDP pairs in each condensate. $\boldsymbol{N}_{\mathbf{C}}\left( \boldsymbol{t} \right)$ continuously undergoes fluctuation, indicating dynamic making and breaking of residue contacts.

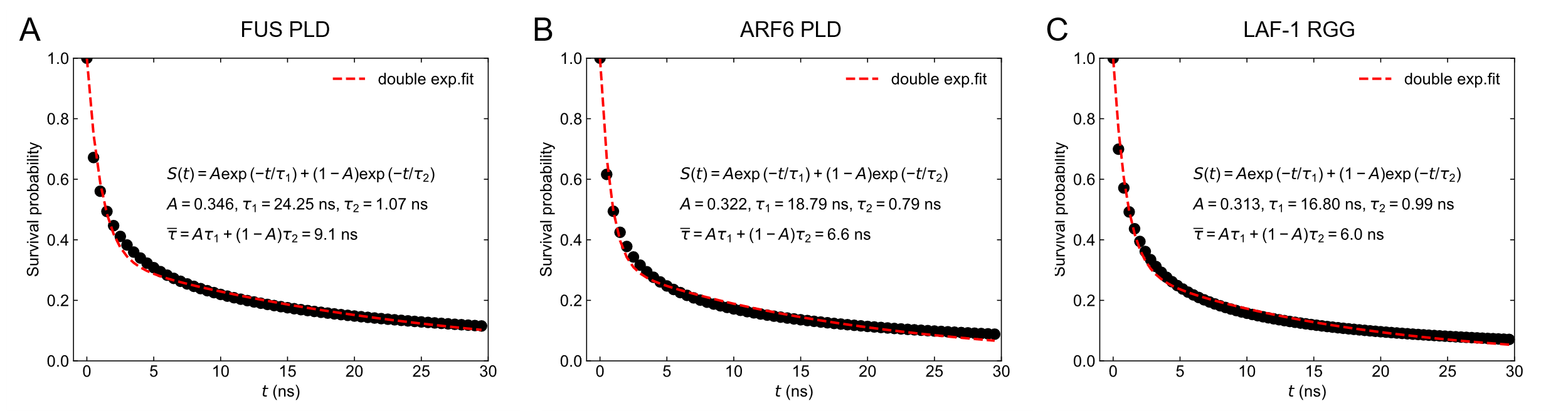
Figure. S6.

Fig. S6. The survival probability of residue contacts *S*(*t*) between IDPs in FUS PLD, ARF6 PLD and LAF-1 RGG condensates. All these survival curves can be described as a double-exponential function (red dashed lines). Three IDP condensate share similar decay modes with $\boldsymbol{\tau}_{\boldsymbol{1}}\boldsymbol{=}$24.25 $\boldsymbol{\pm}$ 0.52 ns, $\boldsymbol{\tau}_{\boldsymbol{2}}\boldsymbol{=}$1.07 $\boldsymbol{\pm}$ 0.02 ns (FUS PLD); $\boldsymbol{\tau}_{\boldsymbol{1}}\boldsymbol{=}$18.79 $\boldsymbol{\pm}$ 0.37 ns, $\boldsymbol{\tau}_{\boldsymbol{2}}\boldsymbol{=}$0.79 $\boldsymbol{\pm}$ 0.02 ns (ARF6 PLD); $\boldsymbol{\tau}_{\boldsymbol{1}}\boldsymbol{=}$16.80 $\boldsymbol{\pm}$ 0.45 ns, $\boldsymbol{\tau}_{\boldsymbol{2}}\boldsymbol{=}$0.99 $\boldsymbol{\pm}$ 0.03 ns (LAF-1 RGG). And the partition coefficient $\boldsymbol{A=}$ 0.346 $\boldsymbol{\pm}$ 0.004 (FUS PLD), $\boldsymbol{A=}$ 0.322 $\boldsymbol{\pm}$ 0.004 (ARF6 PLD), $\boldsymbol{A=}$ 0.313 $\boldsymbol{\pm}$ 0.003 (LAF-1 RGG). The corresponding goodness of fit *R*^2^ are 0.95, 0.97 and 0.98, respectively.

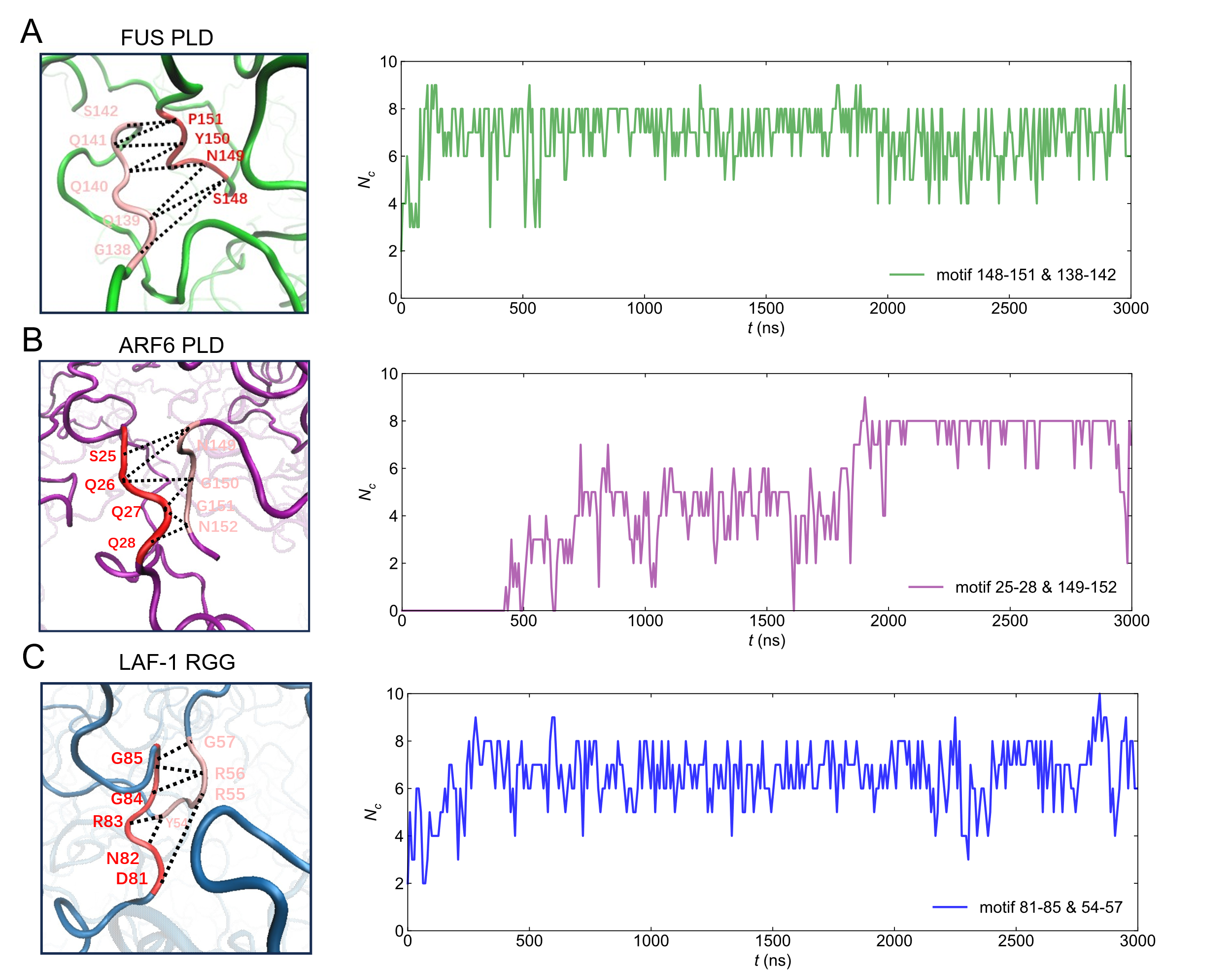
Figure. S7.

Fig. S7. The evolution of the number of residue contacts *N*_c_(*t*) within *motif interaction* in three IDP condensates. (A, B and C) Examples of contact number fluctuations between residue motifs during the 3-μs simulation mentioned in Figure 1C, 1D and 1E, respectively. Notably, the lifetime of these *motif interaction*s all exceed 1000 ns.

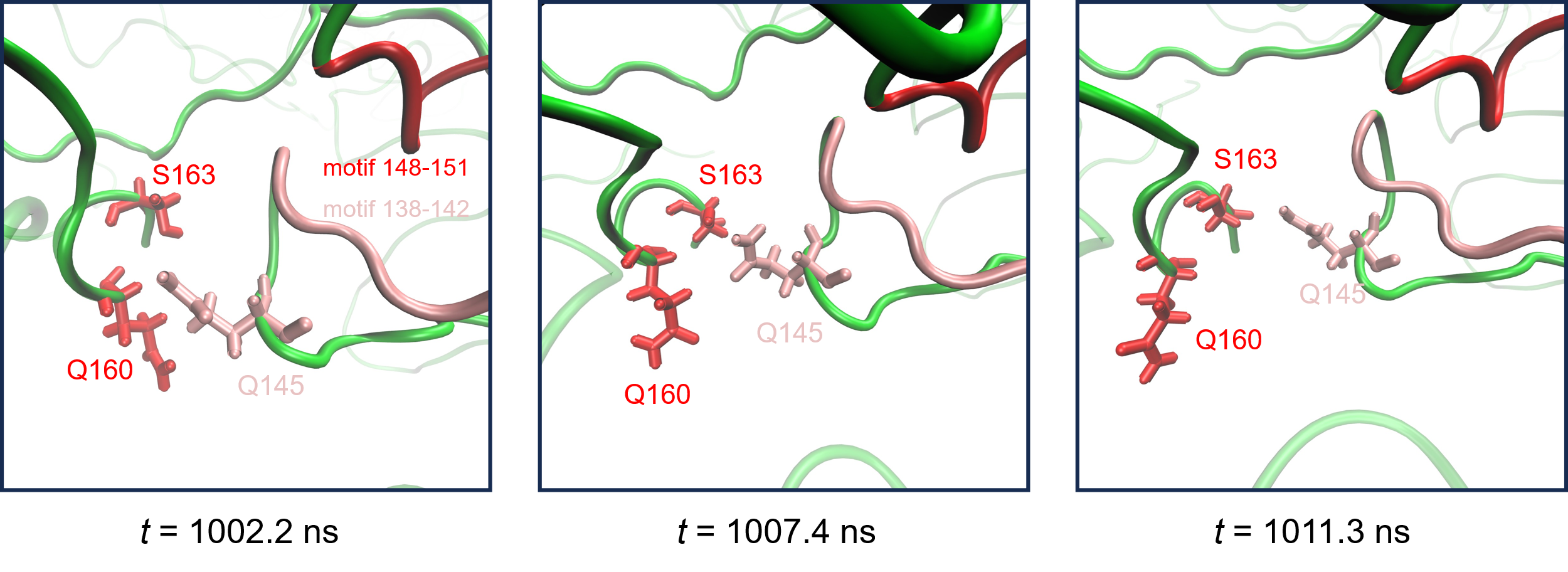
Figure. S8.

Fig. S8. Representative configurations of normal residue contacts in the vicinity of these localized prolong contacts. At *t* = 1002.2 ns, Q145 of IDP-1 contacts with Q160 of IDP-2. This contact quickly breaks at *t* = 1007.4 ns. Subsequently, Q145 can form transient contact with S163 of IDP-2 at *t* = 1011.3 ns.

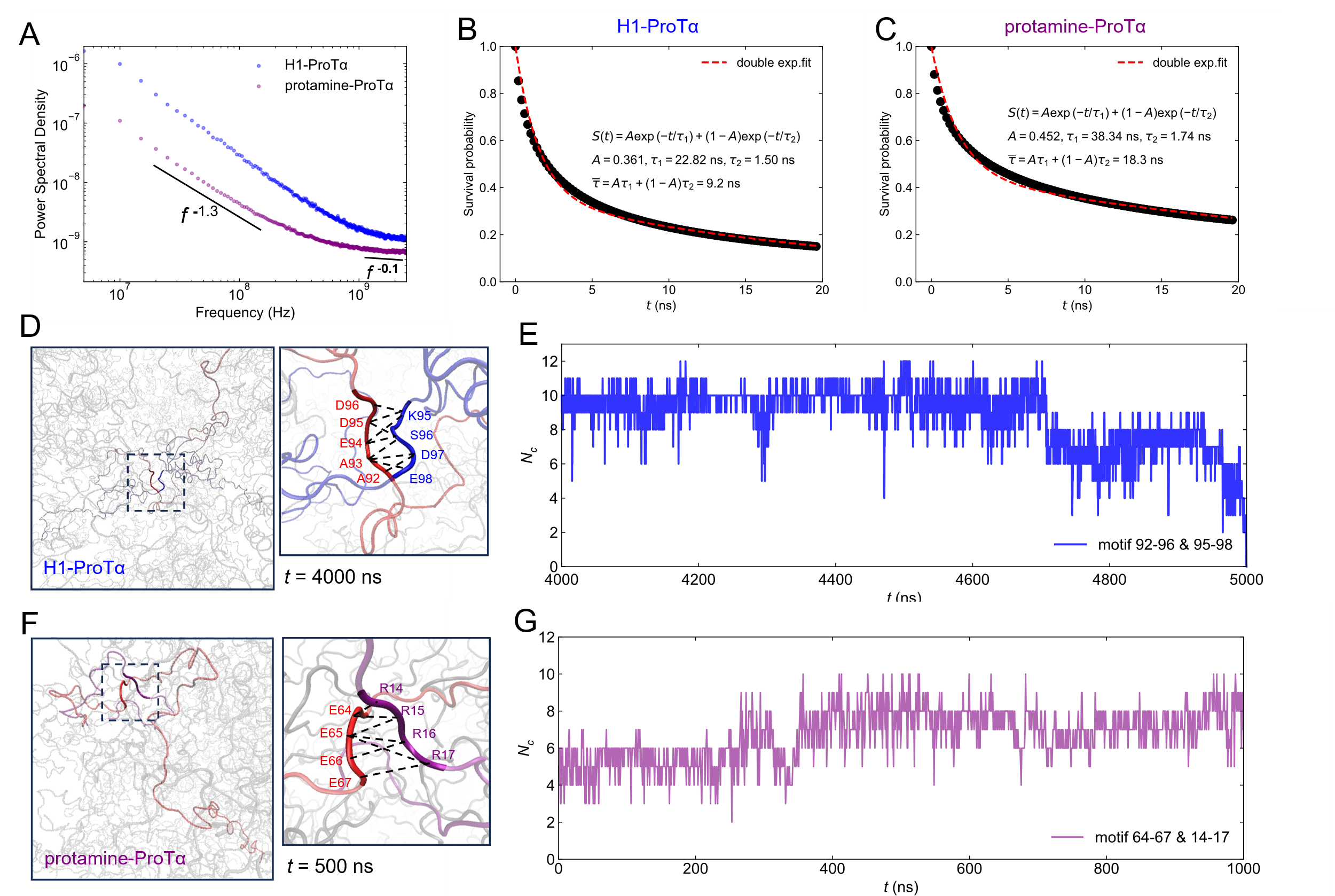
Figure. S9.

Fig. S9. *Motif interaction*s identified in two hybrid condensates. (A) Power spectral density (PSD) of the evolution of the number of residue contacts between IDPs (*N*_c_(*t*)) in H1-ProTα (blue) and protamine-ProTα (purple) condensates. For both systems, the PSD exhibits a plateau above a frequency of ~10^9^ Hz. Notably, at frequencies ~4×10^7^ Hz, PSD exhibits 1/f noise with a power-law exponent of ~-1.3. Hence, there are residue contacts with relatively long lifetimes in these two hybrid condensates. (B, C) The survival probability of residue contacts *S*(*t*) between IDPs. Both the survival curves can be described as a double-exponential function (red dashed lines). The corresponding goodness of fit *R*^2^ are 0.96 and 0.91, respectively. (D) Representative configuration of *motif interaction* within H1-ProTα condensate and localized reorganization of residue interaction. The corresponding residue motifs are 92-96 of chain 117 (red) and 95-98 of chain 123 (blue). And the residue interactions are highlighted in black dashed lines. (E) The contact number fluctuations between motif 92-96 and 95-98. The lifetime of this *motif interaction* exceeds 1000 ns. (F) Representative configuration of *motif interaction* within protamine-ProTα condensate and localized reorganization of residue interaction. The corresponding residue motifs are 64-67 of chain 39 (red) and 14-17 of chain 129 (purple). The residue interactions are highlighted in black dashed lines. (G) The contact number fluctuations between motif 64-67 and 14-17. The lifetime of this *motif interaction* also exceeds 1000 ns.

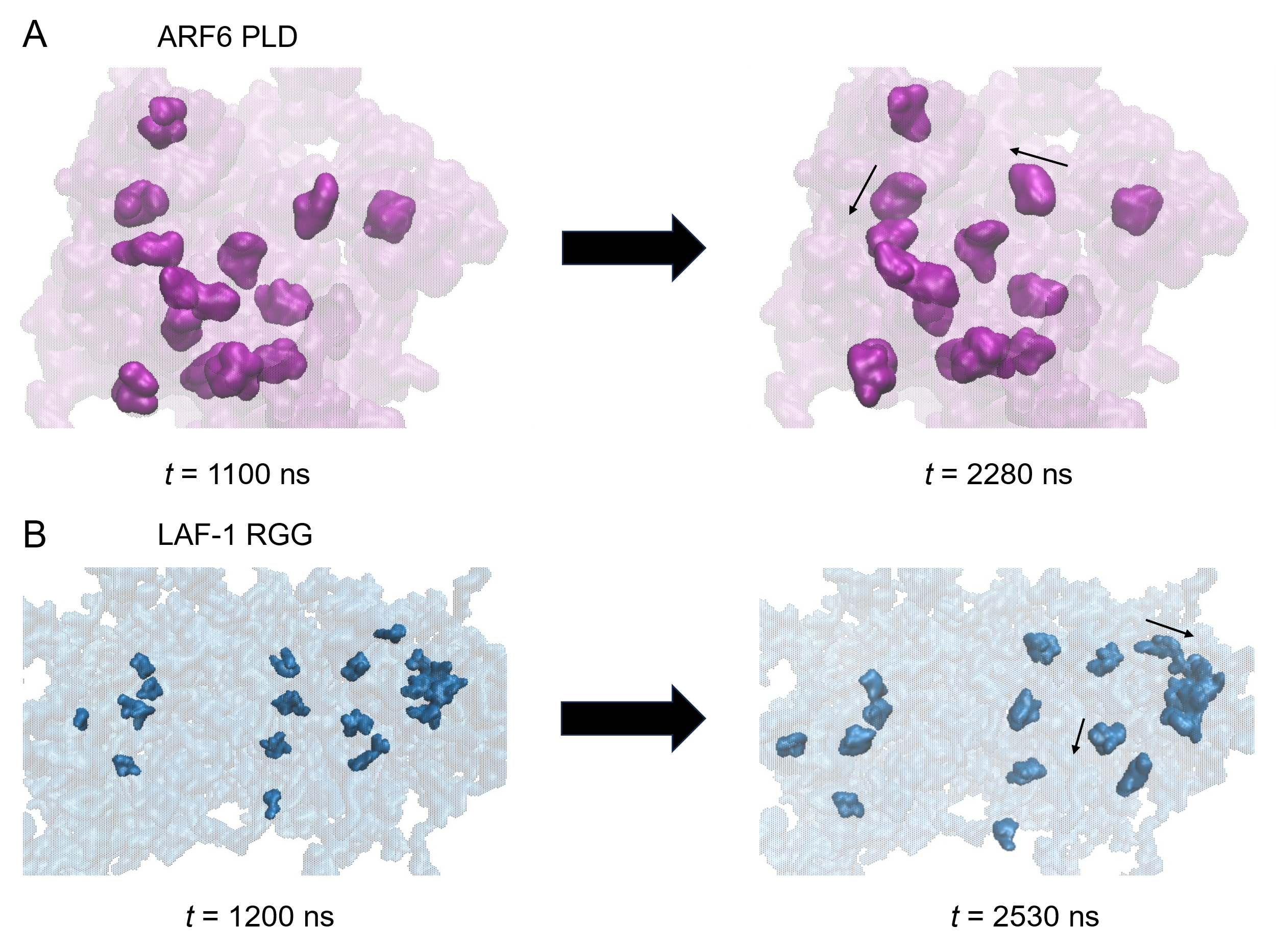
Figure. S10.

Fig. S10. Steady arrangement of HOI-mediated *motif interaction*s within ARF6 PLD and LAF-1 RGG condensates. About 1/2 and 1/3 *motif interaction*s are shown explicitly for ARF6 PLD (A) and LAF-1 RGG (B) condensates, respectively. For both systems, spatial arrangement of *motif interaction*s is quite stable. Only a small fraction of *motif interaction*s undergoes slight slippage (black arrows).

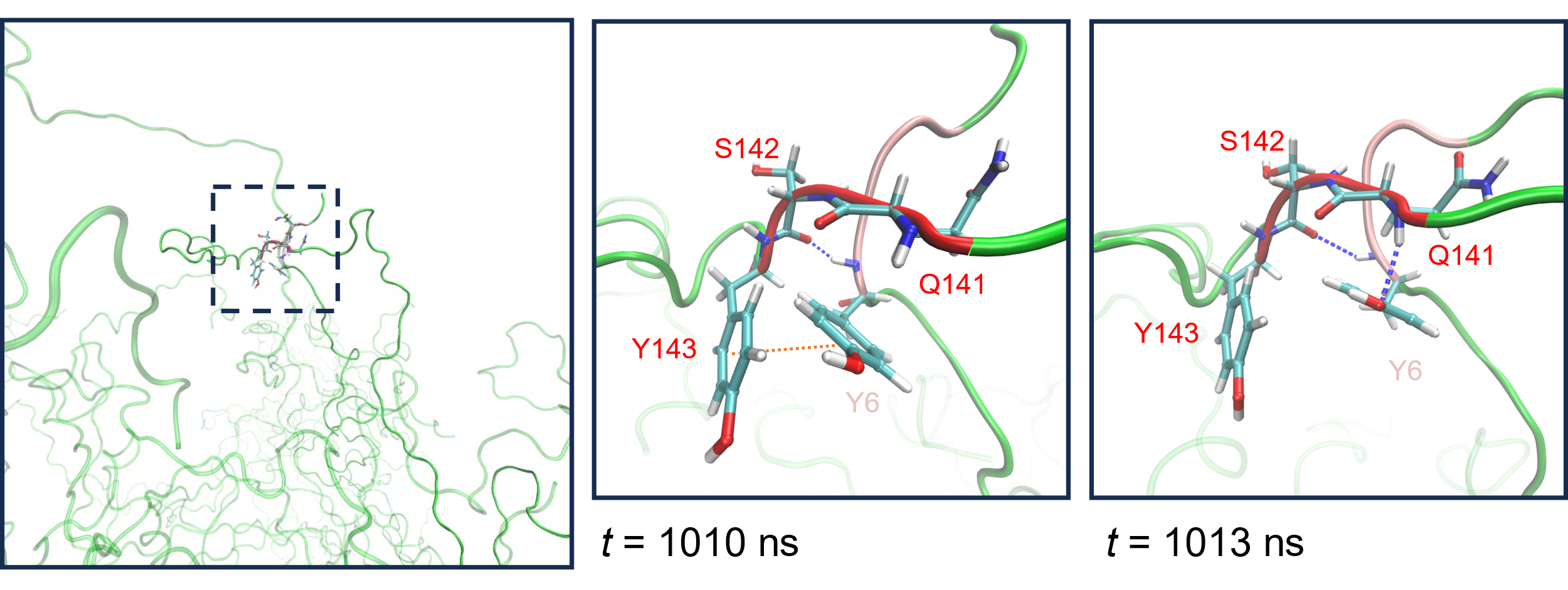
Figure. S11.

Fig. S11. Higher-order residue interactions between HOI-mediated *motif interaction* in FUS PLD condensate. The residue motifs (e.g., 141-143 of chain 2 and 4-6 of chain 3) include 2 tyrosine and 1 glutamine. These residues can interact with multiple residues, and their multivalent interactions undergo local reorganization. For example, at *t* = 1010 ns, Y6 forms hydrogen bond with S142 (blue dashed line) and π-π stacking with Y143 (orange dashed line). At *t* = 1013 ns, the π-π stacking between Y6 and Y143 is substituted by hydrogen bond between Y6 and Q141. Meanwhile, the hydrogen bond between Y6 and S142 remains.

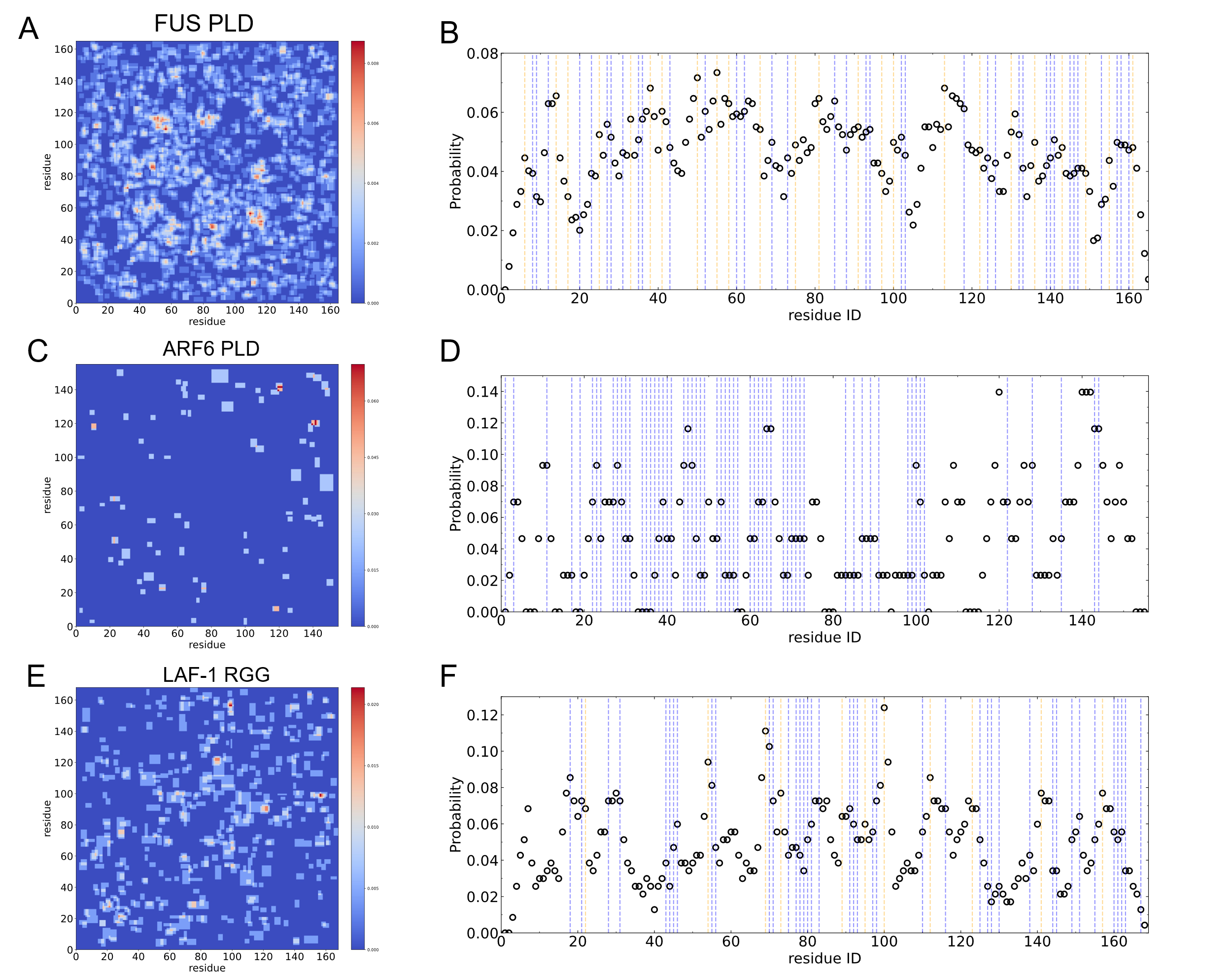
Figure. S12.

Fig. S12. Sequence features of HOI-mediated *motif interaction*s. (A, C and E) Time fractions of the residue pairs participating in *motif interaction* within three condensates. (B, D and F) Time fractions of the residues participating in *motif interaction*. Important residues were highlighted using dashed lines: glutamine (blue dashed lines) and tyrosine (orange dashed lines) of FUS PLD (B) and ARF6 PLD (D), arginine and aspartic acid (blue dashed lines) and tyrosine (orange dashed lines) of LAF-1 RGG (F). Most residues can participate in *motif interaction*, with the probability ranging from 3% to 8%. The average probabilities are 4.7% (FUS PLD), 4.6% (ARF6 PLD) and 4.9% (LAF-1 RGG). For FUS PLD, the probability of tyrosine (5.4%) is slightly higher than other residues. As for LAF-1 RGG, the probability of arginine (5.9%) is mildly above average.

Tables

Table S1. Literature values and references for data shown in Figure S1.

| Liquid condensate | Viscosity (Pa·s) | Interfacial tension (mN/m) |
| --- | --- | --- |
| FIB1 (55) | 100 | 1.23×10^-3^ |
| Cell Nucleus (56) | 3000 | 1.5×10^-3^ |
| Glycinin (57) | ~1600 | ~0.16 |
| NPM1 (*in vivo*) (55) | 37 | 4×10^-4^ |
| Whi3 (58) | 15 | 5×10^-5^ |
| PGL-1 (59) | ~1 | ~10^-3^ |
| Poly R (60) | 14.4 | 0.1 |
| [RGRGG]_5_-dT_40_ (61) | 3 | 0.8 |
| LAF-1 RGG (8, 62) | 10~50 | 0.17 |
| FUS PLD (63-65) | ~4 | ~0.1 |
| ARF6 PLD  (estimated in this work) | 2.3 | 0.48 |
| Silicon oil (66) | 0.02 | 36 |
| Mineral oil (67) | 7.75×10^-3^ | 49 |
| C_16_H_34_ (68) | 2.77×10^-3^ | 55.2 |
| C_10_H_22_ (68) | 9×10^-4^ | 53.2 |
| C_6_H_14_ (68) | 3.13×10^-4^ | 51.4 |
| Water (298K) (64) | 8.9×10^-4^ | 72.8 |
| PAA (69) | ~2×10^-3^ | ~60 |

Table S2. A summary of the setup details of atomistic simulations in this work.

| System | Box size | Number of chains | Simulation time |
| --- | --- | --- | --- |
| FUS PLD | 14×14×35 nm^3^ | 48 | 3×3 μs |
| ARF6 PLD | 13×13×13 nm^3^ | 20 | 3 μs |
| LAF-1 RGG | 10×10×34 nm^3^ | 40 | 3 μs |

Table S3. HOI-mediated *motif interaction*s identified in three homogeneous and two heterogenous condensates.

| System | Total number of interacting IDP pairs | Total number of *motif interaction*s | Average number of *motif interaction*s | Average length of *motif interaction*s |
| --- | --- | --- | --- | --- |
| FUS PLD | 626 | 1053 | 1.68 | 3.79 ± 1.23 |
| ARF6 PLD | 36 | 43 | 1.19 | 3.57 ± 1.25 |
| LAF-1 RGG | 135 | 234 | 1.73 | 4.09 ± 1.66 |
| H1-ProTα^*^  (4 to 5 μs of the trajectory) | 67 | 91 | 1.36 | 4.43 ± 1.57 |
| protamine-ProTα^*^  (128 mM KCl) | 209 | 293 | 1.40 | 4.28 ± 1.49 |

*For heterogenous condensates, i.e., H1-ProTα and protamine-ProTα condensates, *motif interaction*s were identified between different types of IDPs.

Table S4. Interaction energies between IDPs and the contributions of HOI-mediated *motif interaction*s. The average values were taken for all interacting IDP pairs.

| System | Interaction energies  between IDPs (kJ·mol^-1^) | Interaction energies  involving *motif interaction*s(kJ·mol^-1^) |
| --- | --- | --- |
| FUS PLD | -563.7 ± 297.5 | -361.0 ± 231.3 (64.1%) |
| ARF6 PLD | -541.4 ± 416.0 | -295.5 ± 202.2 (54.6%) |
| LAF-1 RGG | -546.8 ± 552.3 | -281.8 ± 440.8 (51.5%) |

Table S5. Data used in calculating various physical properties.

| System | $\tau_{m}$ (ns) | $Rg$(nm) | $\Delta n$ | $V_{m}$(nm^3^) |
| --- | --- | --- | --- | --- |
| FUS PLD | 2196 ± 343 | 4.26 ± 0.48 | 1.54 | ~1.1^**^ |
| ARF6 PLD | 1897± 258 | 2.83 ± 0.79 | ~1^*^ |  |
| LAF-1 RGG | 2421 ± 309 | 3.59 ± 0.69 | 1.22 |  |

*$\Delta n$ of ARF6 PLD cannot be directly determined from the simulation in cubic geometry. According to the values of the other two systems, $\Delta n$ of ARF6 PLD should be close to 1.

**The average volume of amino acids (~0.14 nm^3^ (70)) was used to estimate $V_{m}$ for simplification.

Table S6. Combinations of important residues (i.e., Y, Q, R, D) in HOI-mediated *motif interaction* and their probabilities.

| FUS PLD | | | | | |
| --- | --- | --- | --- | --- | --- |
| Numbers of Y and Q in *motif interaction*s | 1Y  2Q | 1Y  1Q | 2Y  1Q | 1Y  3Q | 2Y  2Q |
| Probabilities | 15.0% | 13.3% | 12.4% | 8.3% | 8.1% |

| ARF6 PLD | | | | | |
| --- | --- | --- | --- | --- | --- |
| Number of Q in *motif interaction*s | 4Q | 2Q | 3Q | 6Q | 5Q |
| Probabilities | 25.6% | 16.3% | 16.3% | 7% | 3.5% |

| LAF-1 RGG | | | | | |
| --- | --- | --- | --- | --- | --- |
| Numbers of Y, R and D in *motif interaction*s | 1R  1D | 2Y  1R  1D | 2R  1D | 1Y  1R  1D | 2Y  1R |
| Probabilities | 6.8% | 5.6% | 5.1% | 5.1% | 5.1% |
